# Locally confined IFN_γ_ production by CD4^+^ T cells provides niches for murine cytomegalovirus replication in the salivary gland

**DOI:** 10.1101/2021.01.14.426650

**Authors:** Josua Oderbolz, Nathan Zangger, Lea Zimmermann, Ioana Sandu, Jörn Starruß, Frederik Graw, Annette Oxenius

## Abstract

Cytomegalovirus (CMV) has evolved a unique virus-host relationship in the salivary glands (SGs) to sustain prolonged viral replication and hence chances for horizontal transmission. Previous reports have established a decisive role for IFN_γ_ producing CD4^+^ T cells to control murine CMV (MCMV) infection in the SGs; however, micro-anatomical information regarding their mode of action is largely missing. Here, we provide a spatiotemporal analysis of defined antiviral immune actions that eventually culminate in control of lytic MCMV replication in this preferred mucosal niche. CXCR3-mediated guidance of CD4^+^ T cells towards CXCL9 and CXCL10 expressing cells resulted in discrete clusters close to infection foci where they reported TCR engagement and produced IFN_γ_. Of note, these clusters occasionally contained CD11c^+^ antigen-presenting cells with engulfed virus-associated remnants, most likely apoptotic bodies derived from previously infected cells, enabling antigen presentation to CD4^+^ T cells. The induced IFN_γ_ production within these CD4^+^ T cell accumulations triggered IFN_γ_R signaling in a confined perimeter, thereby inducing local, but not organ-wide protection, and allowing MCMV replication to continue at not yet protected sites. Combining experimental data with a mathematical model of the spatiotemporal dynamics of infection and CD4^+^ T cell dynamics revealed a scenario, in which ultimate MCMV control is achieved through accumulating sites of regionally-confined tissue protection.

## Introduction

Cytomegalovirus (CMV), a member of the herpesvirus family, is an opportunistic pathogen causing severe clinical outcomes in immunocompromised individuals^1, 2^. Owing to many shared similarities with human CMV (HCMV), including the genetic make-up, the high degree of species specificity and the biological characteristics in their natural habitat such as the huge repertoire of immune evasion mechanisms, infection of mice with murine CMV (MCMV) is a robust infection model to study CMV pathogenesis in an animal setting^3, 4^. Although primary MCMV infection is controlled in most visceral organs within a few days, the salivary glands (SGs) are a peripheral glandular tissue where lytic viral replication is continuing for several weeks^5^. The sustained high viral loads in the SGs facilitate horizontal transmission via saliva. While control of lytic MCMV replication in most tissues is mediated by MCMV-specific CD8^+^ T cells, this is not the case for the SGs^6–8^. MCMV-encoded MHC class I immune evasion genes (i.e. *m04*, *m06* and *m152*) are particularly potent in avoiding recognition of infected cells by cytotoxic CD8^+^ T cells^9, 10^, as deletion of these immune evasion genes restores CD8^+^ T cell recognition of MCMV harboring cells, consequently leading to CD8^+^ T cell-mediated immune control of MCMV infection in the SGs^11^. Therefore, under normal circumstances, control of MCMV infection in the SGs completely relies on CD4^+^ T cells that exert their protective effector functions primarily through the secretion of the pro-inflammatory cytokines interferon gamma (IFN_γ_) and tumor necrosis factor alpha (TNFα)^12, 13^. In this regard, we have previously shown that sensing of CD4^+^ T cell-produced IFN_γ_ by non-hematopoietic cells in the SGs is required for eventual control of lytic viral replication^11^. However, it remains unclear how long-lasting productive virus infection is maintained in this peripheral organ in face of marked infiltration of functional MCMV-specific CD4^+^ T cells early upon infection and the prompt generation of tissue-resident memory T cells^14, 15^. One important aspect that has so far not received much attention is information about micro-anatomical conditions and constraints in the SGs during MCMV infection. This includes spatial information about infection foci, distribution of infiltrating virus-specific CD4^+^ T cells, sites of antigen recognition and IFN_γ_ production, and the range of IFN_γ_ sensing. In the current study, we used advanced microscopy methods to visualize key components of the antiviral immune response with high spatiotemporal resolution. By combining previous knowledge with our experimental data, we further generated a mathematical model that simulates the CD4^+^ T cell-mediated immune control of MCMV infected SGs. We propose a scenario in which MCMV antigens in the SGs are sensed by virus-specific CD4^+^ T cells only in a delayed and indirect manner, after remnants of previously infected cells had been engulfed by local antigen-presenting cells (APCs). This leads to a locally confined IFN_γ_ secretion, affording protection only in this restricted area. However, non-protected areas of the SGs continue to be permissive for infection and replication, evidenced by long-term maintenance of high viral loads in the SGs. Eventual control occurs if local IFN_γ_-concentrations are sufficiently effective to allow agglomeration of protected sites, and thus restriction of viral spread. Thereby, accumulation of virus-specific CD4^+^ T cells by APCs potentiates IFN_γ_ concentrations of otherwise limited IFN_γ_ - release by individual CD4^+^ T cells, which enables prolonged protection of local areas and eventually leading to viral control within the tissue.

## Results

### Virus-specific CD4^+^ T cells infiltrate MCMV-infected SGs and show an activated phenotype

In a first set of experiments, we set out to longitudinally analyze the abundance and phenotype of MCMV-specific CD4^+^ T cells in the SGs upon MCMV infection. As opposed to SGs of naïve mice, which hardly contain any T cells, MCMV infection leads to a substantial infiltration of CD4^+^ and CD8^+^ T cells (**Suppl. Fig. 1a**). To specifically focus on antigen-specific CD4^+^ T cells, we employed T cell receptor (TCR) transgenic (tg) CD4^+^ T cells that specifically recognize the immunodominant MCMV-epitope M25411-425 in the context of MHC class II (referred to as M25 CD4^+^ T cells) in adoptive transfer experiments^16–18^. M25 CD4^+^ T cells were on a CD45.1 background, allowing the distinction from endogenous CD45.2^+^ CD4^+^ T cells (**Suppl. Fig. 1b**). M25 CD4^+^ T cells were adoptively transferred one day prior to infection with recombinant green fluorescent protein (GFP)- expressing MCMV (MCMV-GFP), quantified and phenotypically characterized 8, 14 and 30 days post infection (dpi) in the SGs, spleen and lung (**Fig. 1a**). M25 CD4^+^ T cells expanded and were present in all organs of infected mice, whereas adoptive transfer (AT) of M25 CD4^+^ T cells into naïve or latently MCMV infected mice resulted in low M25 CD4^+^ T cell counts in the SGs (**Suppl. Fig. 1c**). As expected, systemic infection with MCMV-GFP resulted in increasing viral loads in the SGs over time, but not in the spleen and lung where control happened in the first week (**Suppl. Fig. 1d**). We detected the highest percentages and total numbers of M25 CD4^+^ T cells at 8 dpi, followed by a successive reduction over time (**Fig. 1b).** The continuous reduction of SG-resident M25 CD4^+^ T cells from day 8 onwards was accompanied by increasing viral titers and stable numbers of endogenous CD4^+^ T cells (**Suppl. Fig. 1e).** SG-infiltrating M25 CD4^+^ T cells displayed an antigen-experienced CD44^high^CD62L^low^ phenotype and expressed the surface markers LFA-1 (heterodimeric integrin composed of CD11a and CD18) and CD49d, typical features that had been reported previously for MCMV-specific CD4^+^ T cells (**Fig. 1c**)^19–21^. Most of the endogenous CD4^+^ T cells were also CD44^high^LFA-1^+^CD49d^+^, suggesting that the majority of them were also MCMV- specific. Moreover, a considerable fraction of SG-located M25 CD4^+^ T cells expressed the surface markers CD69 and PD-1 8 dpi, associated with an activated status (**Suppl. Fig. 1f).** Interestingly, CXCR3 (being associated with a Th1 phenotype) was almost completely absent in SG- and lung-residing CD4^+^ T cells compared to those in spleen, submandibular lymph node and blood (**Fig. 1d, suppl. Fig. 1g**)^22, 23^.

**Figure 1:**
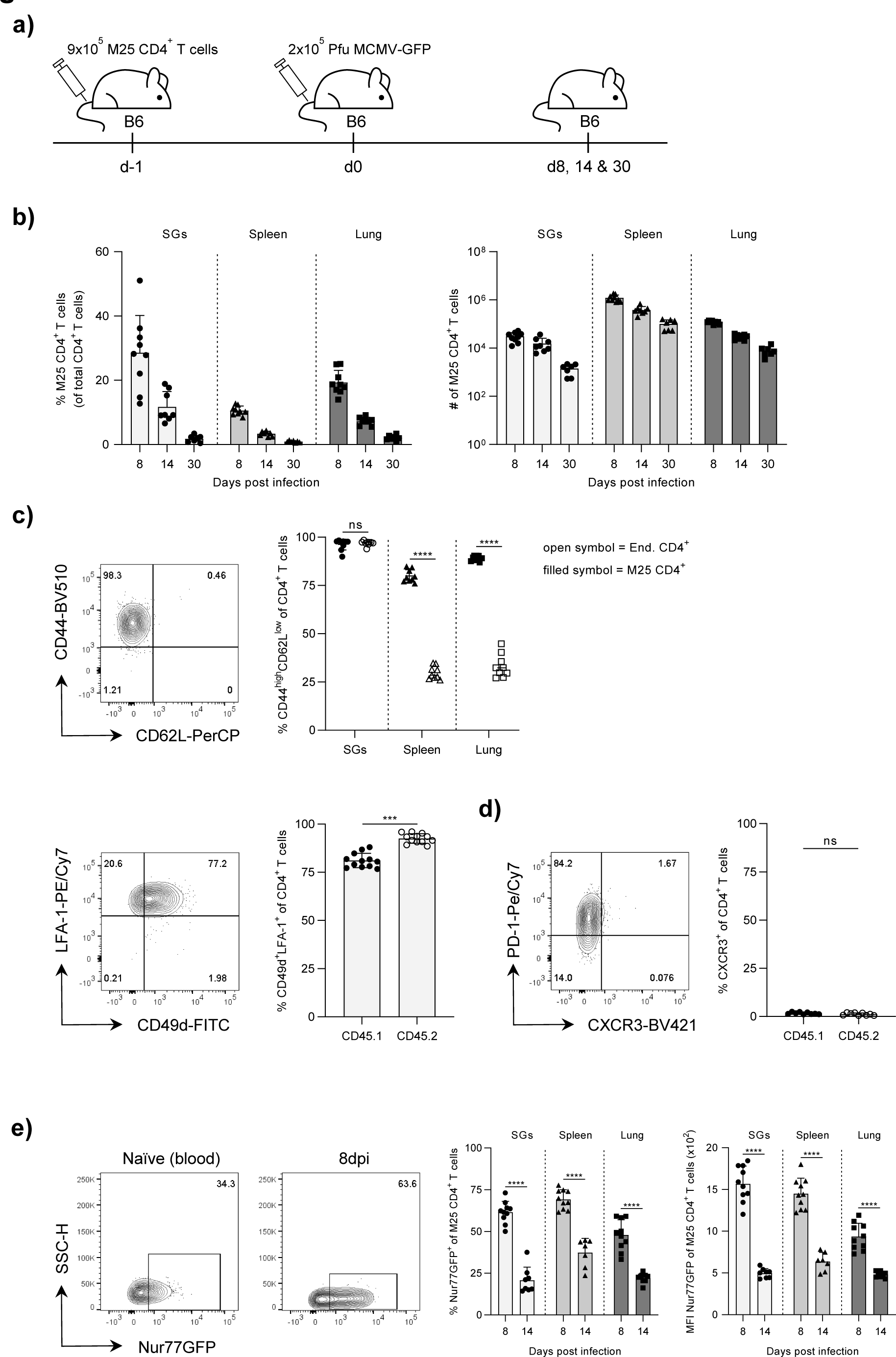
Kinetics and phenotypic characterization of M25 CD4^+^ T cells. **a)** Experimental approach. 9*10^5^ MACS purified M25 CD4^+^ T cells were adoptively transferred into naïve WT B6 mice one day prior MCMV-GFP infection and analyzed in various organs at indicated time points. **b)** Percentage and total number of M25 CD4^+^ T cells in the SGs, spleens & lungs at indicated time points. **c)** Upper row: Representative flow cytometry contour plot of CD44^high^CD62L^low^ M25 CD4^+^ T cells in the SGs (left) and percentage of CD44^high^CD62L^low^ CD4^+^ T cells in the SGs, spleens and lungs (right). Lower row: Representative flow cytometry contour plot of CD49d^+^LFA-1^+^ M25 CD4^+^ T cells (left) and percentage of CD49d^+^LFA-1^+^ CD4^+^ T cells in the SGs (right). **d)** Representative flow cytometry contour plot of CXCR3^+^ M25 CD4^+^ T cells in the SGs (left) and percentage of CXCR3^+^ CD4^+^ T cells in the SGs (right). **e)** Representative flow cytometry contour plot of Nur77GFP^+^ M25 CD4^+^ T cells in the blood of naïve M25xNur77GFP mice and in MCMV-GFP infected SGs (left). Percentage of Nur77GFP^+^ M25 CD4^+^ T cells and MFI of Nur77GFP on M25 CD4^+^ T cells in the SGs, spleens and lungs (right). **c and d)** Representative flow cytometry contour plots and analyses from 8 dpi. Filled symbol = CD45.1^+^ M25 CD4^+^ T cells, open symbol = CD45.2^+^ endogenous CD4^+^ T cells. Data in **b – e** are shown as mean + SD of n = 7 – 12 mice pooled from two independent experiments. Each symbol represents an individual mouse. Statistical significance was determined using single **(c: lower row and d)** or multiple **(c: upper row and e)** unpaired two-tailed t test. *P<0.05, ***P<0.001, ****P<0.0001, ns = not significant.

Next, we asked the question whether M25 CD4^+^ T cells sense their cognate antigen in the SGs. To this end, M25 CD4^+^ T cells harboring a Nur77GFP reporter gene were used (referred to as M25xNur77GFP CD4^+^ T cells) in which TCR engagement via cognate antigen results in GFP expression^24, 25^. M25xNur77GFP CD4^+^ T cells were adoptively transferred one day prior to MCMV-GFP infection and GFP expression in the transferred cells was analyzed 8 and 14 dpi in SGs, spleen and lung. Despite huge differences in viral loads between SGs, spleen and lung, we observed large frequencies of Nur77GFP positive M25 CD4^+^ T cells at 8 dpi with a subsequent decrease at 14 dpi in all organs (**Fig. 1e, suppl. Fig. 1d).** These data suggest that viral loads are not a good proxy for the amount of antigen sensed by M25-specific CD4^+^ T cells. Similar results were obtained when a recombinant mcherry-expressing MCMV strain (MCMV-3DR) was used for the infection, which caused higher viral loads in the SGs within the first two weeks after infection compared to the MCMV-GFP mutant (**Suppl. Fig. 1h and i).** Taken together, these results indicate that SG-infiltrating M25 CD4^+^ T cells have an activated, antigen-experienced phenotype with substantial exposure to cognate antigen during early stage of MCMV infection in the SGs.

### *In vivo* M25 peptide challenge leads to pronounced IFNγ production by SG-localized M25 CD4^+^ T cells in a TCR-dependent manner

Previous studies investigated the pro-inflammatory cytokine secretion profile of MCMV-specific CD4^+^ T cells upon *ex vivo* exposure to a defined pool of peptides or lysate of MCMV-infected cells^16, 18, 19^. To evaluate the potential of SG-residing M25 CD4^+^ T cells to produce the pro-inflammatory cytokines IFN_γ_ and TNFα, we systemically challenged mice that had received M25 CD4^+^ T cells with the cognate M25 peptide at day 8 or 21 post MCMV infection and harvested the SGs and spleen three hours later (**Fig. 2a**)^20^. The majority of M25 CD4^+^ T cells in both organs upregulated the early T cell activation marker CD69 upon *in vivo* M25 peptide administration compared to the DMSO control injection (**Fig. 2b).** Moreover, a considerable proportion of M25 CD4^+^ T cell expressed IFN_γ_ and a surprisingly low percentage co-expressed IFN_γ_ and TNFα in the SGs (**Fig. 2c**). Notably, comparable frequencies of M25 CD4^+^ T cells upregulated CD69 and produced IFN_γ_ upon *in vivo* M25 peptide challenge at 8 and 21 dpi (**Fig. 2d).** Next, we validated that IFN_γ_ production was due to TCR engagement. We used M25xNur77GFP CD4^+^ T cells for the AT, followed by MCMV infection and M25 peptide challenge at 8 dpi. Over 90% of M25 CD4^+^ T cells in the SGs and spleen turned Nur77GFP positive upon M25 peptide administration (**Fig. 2e).** Furthermore, IFN_γ_^+^ cells were all Nur77GFP^+^, revealing that indeed the IFN_γ_ production is the result of TCR-based antigen recognition and downstream signaling (**Fig. 2f**). Of note, the reduced percentage of Nur77GFP^+^ M25 CD4^+^ T cells at day 8 post MCMV infection in the SGs and spleen compared to results shown in Figure 1 (**Fig. 1e and suppl. Fig. 1i**) is due to the paraformaldehyde (PFA)-based fixation procedure. These data show that M25 CD4^+^ T cells can be locally triggered in the SGs after systemic administration of the cognate peptide, and that 10 - 40% of M25 CD4^+^ T cells produce IFN_γ_ in a TCR-dependent manner.

**Figure 2:**
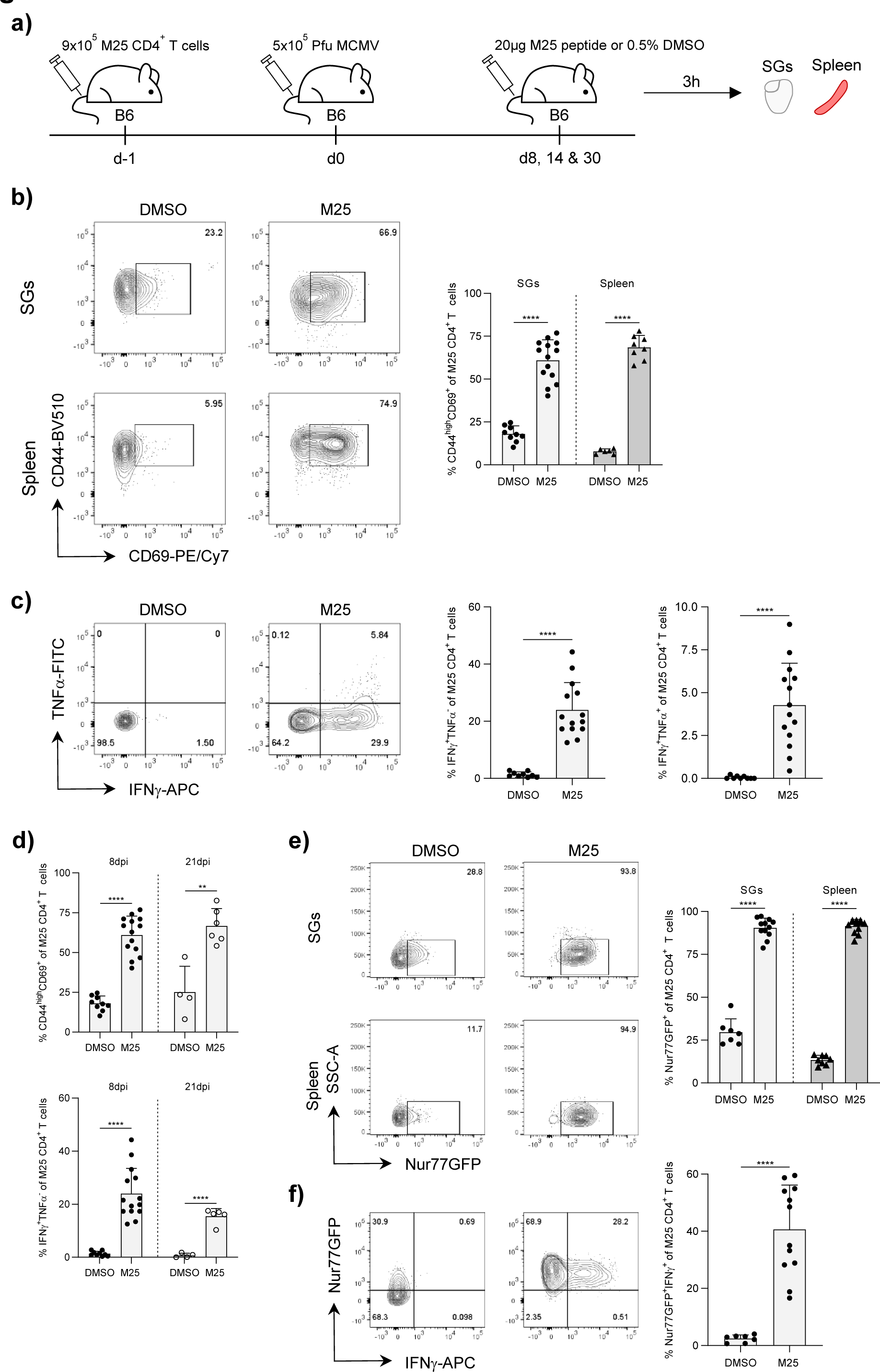
IFN*γ* production by SG-localized M25 CD4^+^ T cells. **a)** Experimental approach. 10^6^ MACS purified M25 (Nur77GFP) CD4^+^ T cells were adoptively transferred into naïve WT B6 mice one day prior MCMV infection and subsequently challenged for 3h with i.v. administered cognate M25 peptide or 0.5% DMSO as control at day 8 or 21 post MCMV infection. **b)** Representative flow cytometry contour plots (left) and quantification (right) of early activated CD44^high^CD69^+^ M25 CD4^+^ T cells in the SGs and spleens 3h post M25 peptide or DMSO administration at day 8 post MCMV infection. **c)** Representative flow cytometry contour plots (left) and quantification (right) of IFN_γ_ and TNFα expression levels by M25 CD4^+^ T cells in the SGs 3h post M25 peptide or DMSO administration at day 8 post MCMV infection. **d)** Percentage of CD44^high^CD69^+^ (top) and IFN_γ_^+^ (bottom) M25 CD4^+^ T cells 3h post M25 peptide or DMSO administration at day 8 and 21 post MCMV infection. **e)** Representative flow cytometry contour plots (left) and quantification (right) of Nur77GFP^+^ M25 CD4^+^ T cells in the SGs and spleens 3h post M25 peptide or DMSO challenge at day 8 post MCMV infection. **f)** Representative flow cytometry contour plots (left) and quantification (right) of Nur77GFP^+^IFN_γ_^+^ M25 CD4^+^ T cells in the SGs 3h post M25 or DMSO challenge at day 8 post MCVM infection. Data in **b – f** are shown as mean + SD of n = 4 – 14 mice pooled from two (b: spleen and d: 21dpi) or three independent experiments. Each symbol represents an individual mouse. Statistical significance was determined using single **(c and f)** or multiple **(b, d and e)** unpaired two-tailed t test. **P<0.01, ****P<0.0001.

### Entire cross section analysis of acutely MCMV-infected SGs reveals micro-anatomical localization of early infiltrating CD4^+^ T cells

Next, we studied the micro-anatomical localization of M25 and endogenous CD4^+^ T cells in acutely MCMV infected SGs by adding another layer of information and resolution. For this purpose, we transferred traceable RFP-expressing M25 CD4^+^ T cells (referred to as M25xRFP CD4^+^ T cells) and subsequently infected mice with MCMV-GFP. Eight days post MCMV-GFP infection, we harvested the SGs and performed whole slide imaging (WSI), followed by in-depth spatial relation studies between SG-infiltrating CD4^+^ T cells and large actively infected (GFP^+^) cells (sites of infectious virus production) using the quantitative image analysis software HALO (**Fig. 3a).** To prevent tissue damage, we carefully dissected, cut and imaged the tightly connected submandibular -and sublingual glands (SMGs and SLGs, respectively) together. Differences in structural features enabled the separation of the SGs into SMGs and SLGs (**Suppl. Fig. 2a).** By applying four-color entire cross section (ECS) analyses, we simultaneously visualized endogenous CD4^+^ and M25 CD4^+^ T cells, along with sites of viral replication. M25 CD4^+^ T cells were readily identified by their co-expression of cytoplasmic RFP and the cell membrane CD4 marker, whereas endogenous CD4^+^ T cells were only positive for the latter. Sites of viral replication were identified by their “Owl’s eye” appearance of well-known inclusion bodies (enlarged cells that are bright for the virus-encoded fluorescent reporter) (**Fig. 3b**)^26^. First, we analyzed total numbers of M25 CD4^+^ T cells in the SMGs and SLGs and detected the majority of M25 CD4^+^ T cells in the SMGs (**Fig. 3c**). However, this was not surprising since the SMGs encompass an approximately six times larger surface area than the SLGs (**Suppl. Fig. 2b**). Nevertheless, normalizing for the surface area also showed a higher density of M25 CD4^+^ T cells in the SMGs (**Fig. 3c**). Next, we quantified the sites of viral replication across the sections and observed considerable heterogeneity, ranging from one to twelve sites of viral replication per SMG cross section (**Fig. 3d**). As CD4^+^ T cell density and number of infected cells were higher in the SMGs, we focused the ensuing spatial analyses to this part of the SGs. In this regard, we quantified distances between sites of viral replication and endogenous or M25 CD4^+^ T cells by extracting data of interest from raw images and employing nearest neighbor analysis. Based on the distribution of measured distances, we quantified the frequencies of endogenous CD4^+^ and M25 CD4^+^ T cells residing within a defined radius from the next site of viral replication (**Suppl. Fig. 2c).** Proximity analyses revealed that most of the endogenous CD4^+^ and M25 CD4^+^ T cells were located 600 μm and more away from the next site of viral replication. However, the percentage of M25 CD4^+^ T cells in relative close proximity to these infection foci (≤ 300 μm) was significantly higher than the ones of the endogenous CD4^+^ T cell compartment (**Fig. 3e).** Given the considerable differences in numbers of viral replication sites per SMG section and the rather random distribution of M25 CD4^+^ T cells at 8 dpi, we expected a correlation between the proportion of M25 CD4^+^ T cells residing within a radius of 300 μm to the nearest site of viral replication and the total counts of sites of viral replication per ECS. Indeed, we discovered a tendency that the number of infection foci strongly influenced this frequency (**Fig. 3f**). However, a few tissue sections showed signs of virus-associated T cell clustering, noticeable by the high frequency of M25 CD4^+^ T cells within 300 μm to a relatively low number of sites of viral replication (**Fig. 3f & g).** We further quantified the Euclidean distance between two closest endogenous CD4^+^ and M25 CD4^+^ T cells and observed slightly shorter distances between M25^+^ CD4^+^ T cells compared to the endogenous CD4^+^ T cells (**Fig. 3e**). Generally, both CD4^+^ T cell populations appeared to be distributed quite randomly and rarely showed signs of accumulation close to infection foci. Therefore, the variation in the average distances between two M25 CD4^+^ T cells is mostly due to differences in cell densities, thus indicating that the total number of M25 CD4^+^ T cells per tissue section has a stronger impact on the distances between these cells than distinct accumulation characteristics (**Fig. 3h).** Finally, *in vivo* short-term intravenous (i.v.) antibody labeling revealed that M25 CD4^+^ T cells do not reside in close proximity to the circulation (**Suppl. Fig. 2d).** In summary, these results indicate that M25 and endogenous CD4^+^ T cells are not preferentially found in close proximity to sites of infection, but rather distribute randomly across the tissue of acutely MCMV-GFP infected SGs.

**Figure 3.**
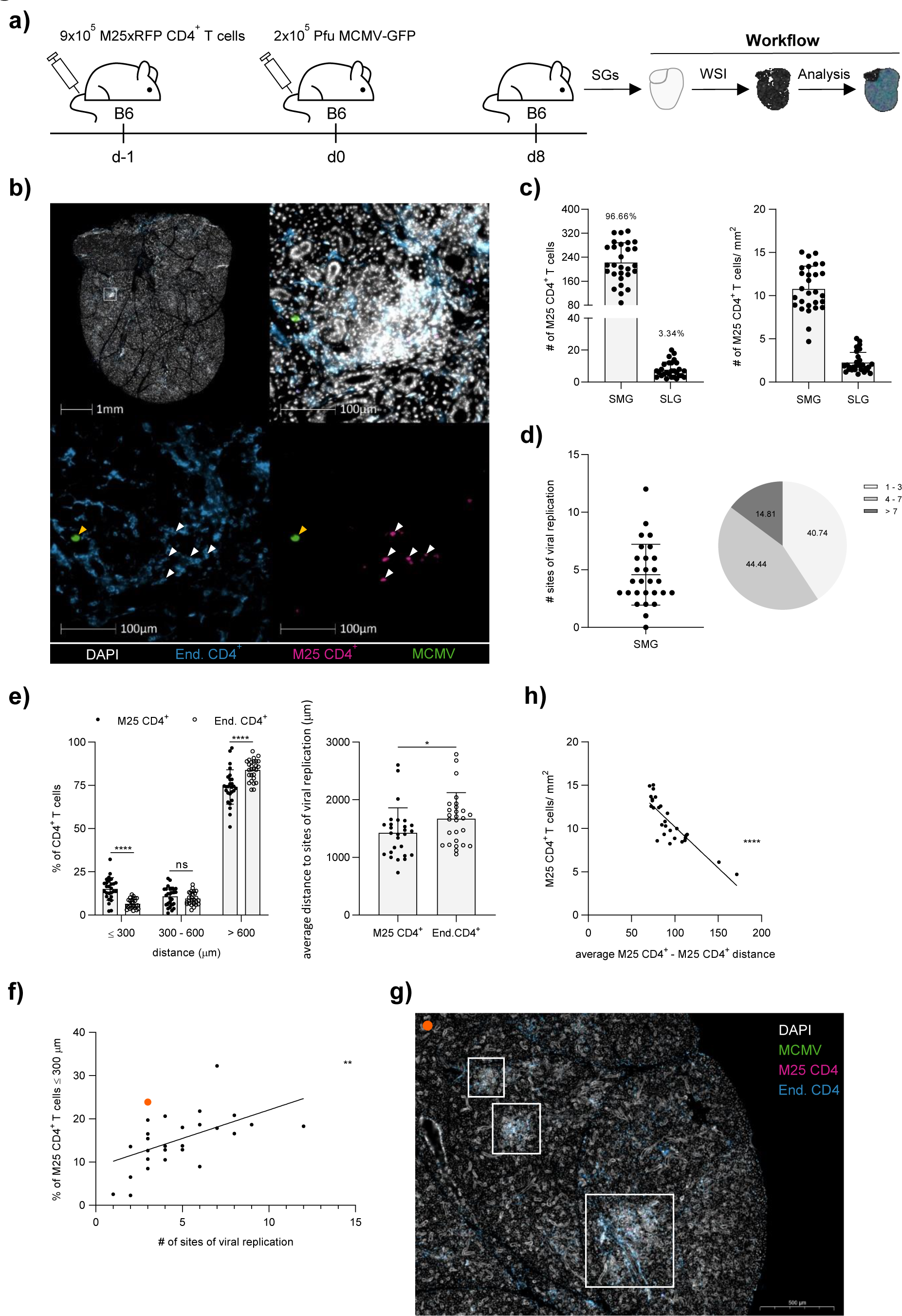
Spatial localization of endogenous CD4^+^ and M25 CD4^+^ T cells in acutely MCMV infected SGs. **a)** Experimental approach. 9*10^5^ MACS purified RFP traceable M25 CD4^+^ T cells were adoptively transferred into naïve WT B6 mice one day prior MCMV-GFP infection. 8 days post MCMV-GFP infection, 10μm thin tissue sections of the SGs were sliced and subjected to whole slide imaging (WSI), followed by quantitative image analysis using the HALO^®^ software. **b)** Example of an entire cross section (ECS) of the right SG lobe (upper left) with indicated magnified region (upper right). Endogenous CD4^+^ T cells are shown in cyan (bottom left), M25 CD4^+^ T cells are shown in magenta (bottom right, white arrowheads), and a site of viral replication is shown in green (bottom, yellow arrowhead). **c)** Total M25 CD4^+^ T cell counts (left) and cell density (right) in the SMG and SLG. **d)** Number of sites of viral replications in the SMG (left) with the percental distribution (pie chart, right). **e)** Percentage of endogenous CD4^+^ and M25 CD4^+^ T cells located within a defined distance to the nearest infection foci (left) and summarized average distances of the endogenous CD4^+^ and M25 CD4^+^ T cells to these sites (right). **f)** Correlation between the number of infection foci per ECS and the percentages of M25 CD4^+^ T cells located within 300 μm away from the nearest infection foci. **g)** Cutout of an ECS of a B6 mouse-derived SG 8 days post MCMV-GFP infection. White frames highlight sites of T cell accumulations close to sites of active viral replication. Data point represented by the orange dot in **f)** corresponds to the cutout of the ECS in **g). h)** Correlation between the cell densities of M25 CD4^+^ T cells and the average in between distances per ECS. Data in **c – f & h** are shown as mean + SD of n = 5 mice (2 -11 ECS per SG, total 28 ECS) pooled from two independent experiments. Each dot represents an individual ECS. Total 275’359 endogenous CD4^+^ and 6’218 M25 CD4^+^ T cells were analyzed. Statistical significance was determined using single (**e, right**) multiple (**e, left**) unpaired two-tailed t test, or two-tailed Pearson correlation (**f & h**). **P<0.01, ****P<0.0001, ns = not significant. Scale bar in **b** = 100 μm and in **g** = 500 μm. SMG = Submandibular gland, SLG = Sublingual gland.

### CD4^+^ T cells represent the major cellular source of IFN*γ* production in MCMV infected SGs

The pro-inflammatory cytokine IFN_γ_ was previously shown to be a crucial component of the antiviral control mechanism in MCMV infected SGs^12^. Therefore, we evaluated next which immune cell type present in the MCMV-infected SGs would preferentially produce the pivotal cytokine IFN_γ_. To this end, we made use of the IFN_γ_ reporter mouse GREAT, which expresses the enhanced yellow fluorescence protein (EYFP) under the control of the IFN_γ_ promoter^27^. In doing so, we investigated cell-specific and spatial IFN_γ_ production in MCMV-3DR infected SGs. We infected GREAT mice with MCMV-3DR and determined which cell types were EYFP^+^ (i.e. IFN_γ_ producing) at 14 dpi. Compared to naïve GREAT mice, we observed significantly elevated EYFP signal levels upon MCMV-3DR infection (**Fig. 4a, suppl. Fig. 3a).** We then characterized the cell types within the EYFP positive cell fraction and identified CD3^+^ cells as the primary cellular source of IFN_γ_ production in MCMV infected SGs. A minor proportion was assigned to NK1.1^+^ natural killer (NK) cells (**Fig. 4b, suppl. 3b)**. Further classification of the EYFP^+^CD3^+^ cell population revealed that ¾ of these cells were CD4^+^ T cells (**Fig. 4c, suppl. Fig. 3c).** Additionally, we measured EYFP signal intensities in total CD4^+^, CD8^+^ and NK1.1^+^ cells. Again, CD4^+^ T cells showed the highest percentage of EYFP^+^ cells, followed by NK cells and CD8^+^ T cells (**Fig. 4d, suppl. Fig. 3d).** Next, we aimed to study IFN_γ_ producing cells *in situ* by applying confocal microscopy. We therefore performed microscopic analyses of MCMV-3DR infected SGs to reveal spatial relations between IFN_γ_-producing CD4^+^ and CD8^+^ T cells and sites of viral replication. We simultaneously recorded CD4^+^, CD8^+^ and EYFP^+^ cells in various field of views (FOVs) of MCMV-3DR infected SGs containing at least one site of viral replication, and attributed each EYFP^+^ cell to its corresponding cell type (**Fig. 4e & f).** In line with the flow cytometry analyses, CD4^+^ T cells were the major cellular source of IFN_γ_ production (**Fig. 4g**). Moreover, spatial analyses revealed that CD4^+^ T cells tended to accumulate in dense clusters at 14 dpi, whereas CD8^+^ T cells showed a more dispersed distribution, despite higher abundance (**Suppl. Fig. 3e**). Of note, additional microscopic analyses of MCMV-3DR-infected SGs from Nur77GFP mice further identified dense CD4^+^ T cell clusters in vicinity to infectious virus production. This implies that accumulations of CD4^+^ T cells close to sites of viral replication represent primary sites of antigen recognition, identifiable by a remarkable fraction of GFP expressing CD4^+^ T cells. (**Fig. 4h & suppl. Fig. 3f & g)**. Interestingly, we sometimes detected small structures containing the mcherry signal from the MCMV-3DR strain within these CD4^+^ T cell accumulations, congruent with a high abundance of mainly EYFP^+^ IFN_γ_ producing CD4^+^ T cells (**Fig. 4i).** In this regard, further microscopic analyses identified CD11c^+^ APC as primary cell type harboring these small mcherry^+^ remnants of MCMV-infected cells, whereas active lytic viral replication was predominantly found within E-Cadherin^+^ epithelial cells (**Suppl. Fig. 4a & b)**. These results strongly confirm previous findings from our lab, that the presentation of phagocytosed remnants (presumably apoptotic bodies containing viral proteins) from previously infected cells by CD11c^+^ APCs lead to local CD4^+^ T cell activation^11^. Thus, CD4^+^ T cells rather recognize apoptotic cell-derived viral structures presented by APCs than directly infected epithelial cells. Taken together, these findings show that CD4^+^ T cells are the major cellular source of IFN_γ_ production in MCMV-infected SGs, and that IFN_γ_-producing CD4^+^ T cells are found in dense clusters at 14 dpi, usually in close proximity to sites of active viral replication. Furthermore, local TCR triggering and IFN_γ_ production is often associated with the presence of CD11c^+^ cells containing mcherry^+^ inclusions - most likely remnants from infected cells, serving as antigen source for the activation of CD4^+^ T cells.

**Figure 4.**
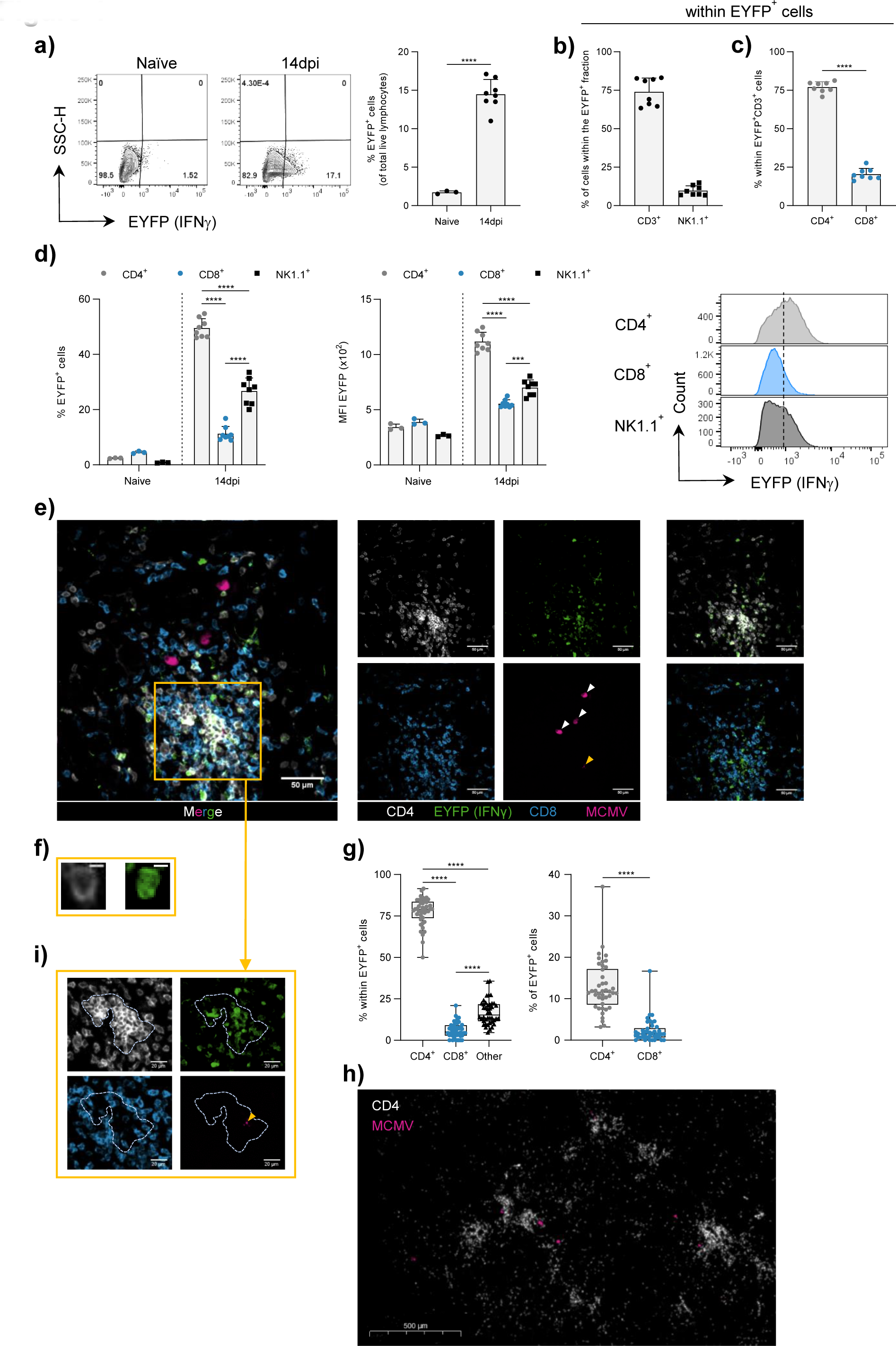
IFN*γ* production by CD4^+^ T cells in MCMV infected SGs. **a)** Representative flow cytometry contour plots of naïve and MCMV-3DR infected Great mice (left) and percentage of total EYFP^+^ cells (right), **b)** percentage of CD3^+^ and NK1.1^+^ cells within EYFP^+^ cells, and **c)** percentage of CD4^+^ and CD8^+^ T cells within CD3^+^EYFP^+^ cells. **d)** Percentage of EYFP^+^ cells within the indicated cell types (left) and the MFI of the EYFP signal (middle). Representative histograms of the EYFP expression profile in the indicated cell types 14dpi (right). Dashed line defines threshold for positive signal. **e)** Example of a four color FOV of a Great mouse-derived SG 14 days post MCMV-3DR infection (left). Single color channel images (middle), merged two color CD4/EYFP (right, upper row) and CD8/EYFP (right, lower row) images. **f)** Magnified IFN_γ_^+^ CD4^+^ T cell (cytosolic EYFP and cell membrane CD4 signal). **g)** Quantification of microscopic analyses. Percentage of cell types within total counted EYFP^+^ cells (left) and the frequency of EYFP^+^ cells within CD4^+^ and CD8^+^ T cells (right) per FOV of MCMV-3DR infected SGs 14dpi. **h)** Cutout of an ECS of a Nur77GFP mouse-derived SG 14 days post MCMV-3DR infection. **i)** Magnified region split in four single color channel images. Dashed line demarcates dense accumulation of preferentially IFN_γ_^+^ CD4^+^ T cells in close proximity to faint mcherrry signal (yellow arrowhead). Data in **a – d** and are shown as mean ± SD of n = 3 naïve and n = 8 infected Great mice pooled from 2 independent experiments. Data in **g** are shown as box and whiskers showing all points from Min. to Max. (n = 42 FOVs with n > 6 per mice) of n = 3 mice from one independent experiment. Each symbol in **a – d** represents one individual mouse and in **g** one individual FOV. Statistical significance was determined using single unpaired two-tailed t test **(a – c** and **g: right**), or 2-way Anova with post hoc Tukey’s multiple comparisons test **(d and g: left)**. ***P<0.001, ****P<0.0001. Scale bar in **e** = 50 μm, in **f** = 5 μm, in **h** = 100 μm and in **i** = 20 μm. White and yellow arrowheads in **e** and **i** indicate infection foci and presumably cargo/ remnants of infected cells, respectively. FOV = Field of view.

### 3D confocal imaging provides a new level of information regarding the CD4^+^ T cell - MCMV interaction

Classical FOV analyses along with the WSI approach allowed us to investigate antiviral immune responses in MCMV infected SGs in two dimensions; however, organs are intrinsically built up in three dimensions. Moreover, the considerable heterogeneity of number of sites of viral replication, together with their anatomical location and complex interrelation with CD4^+^ T cells, required a more profound three-dimensional (3D) examination of the tissue sample. Furthermore, dense clusters of CD4^+^ T cells surrounding virus-associated small structures needed to be confirmed and explored in more detail. For this purpose, we elaborated a clearing-based 3D confocal microscopy pipeline for 200 μm thick tissue sections (**Fig. 5a**) ^28, 29^. In doing so, PFA fixed MCMV-3DR infected SGs were first embedded in agarose and subsequently sliced at appropriate thickness using a vibrating microtome. Tissue sections were then stained, optically cleared, and imaged using an inverted confocal microscope. Finally, 3D reconstructions composed of sequentially acquired 2D-stacked images were processed and visualized, followed by in-depth spatial analyses. As proof, tissue clearing with the histodenz-based refractive index (RI) matching solution (RIMS) substantially increased imaging depth up to 200 μm, allowing the precise identification of several sites of viral replication and the truthful evaluation of CD4^+^ T cell counts (**Fig. 5b & Suppl. Video 1).** Furthermore, and in line with observations from the 2D images, specific FOVs at different z-planes revealed the emergence and disappearance of accumulating CD4^+^ T cells around small virus-associated structures, whereas closely situated large infection foci seemed to be ignored (**Fig. 5c).** Nearest neighbor analysis further confirmed the observation that a relevant number of CD4^+^ T cells are in close proximity to many of these small mcherry^+^ remnants, which presumably represent apoptotic bodies released from previously infected cells (**Fig. 5d, Suppl. Fig. 4c & Video 2).** Image segmentation of actively MCMV-3DR-infected cells and small virus-associated structures derived from infected cells revealed different morphological appearances (**Fig. 5e & suppl. Fig. 4c**). As previously indicated (**Suppl. Fig. 4a & b),** small mcherry^+^ virus-associated remnants were frequently found within CD11c^+^ APCs (≈ 30%), most likely as a result of phagocytic activity **(Suppl. Video 3)**. In contrast, large actively infected cells were only assigned to E-Cadherin^+^ epithelial cells, again confirming previous knowledge that acinar glandular epithelial cells represent the preferred cell type in the SGs for infectious virus production (**Fig. 5f & g)**. Importantly, unassigned mcherry signals of small volumes might be cell-free (not yet phagocytosed) or found within other cell types, for instance in CD11b^+^ cells (**Suppl. Fig. 4d**) In summary, 3D confocal imaging confirmed that dense accumulations of CD4^+^ T cells are often associated with APCs that harbor cargo from previously virus-infected cells, likely remnants or apoptotic bodies **(Suppl. Video 4)**.

**Figure 5.**
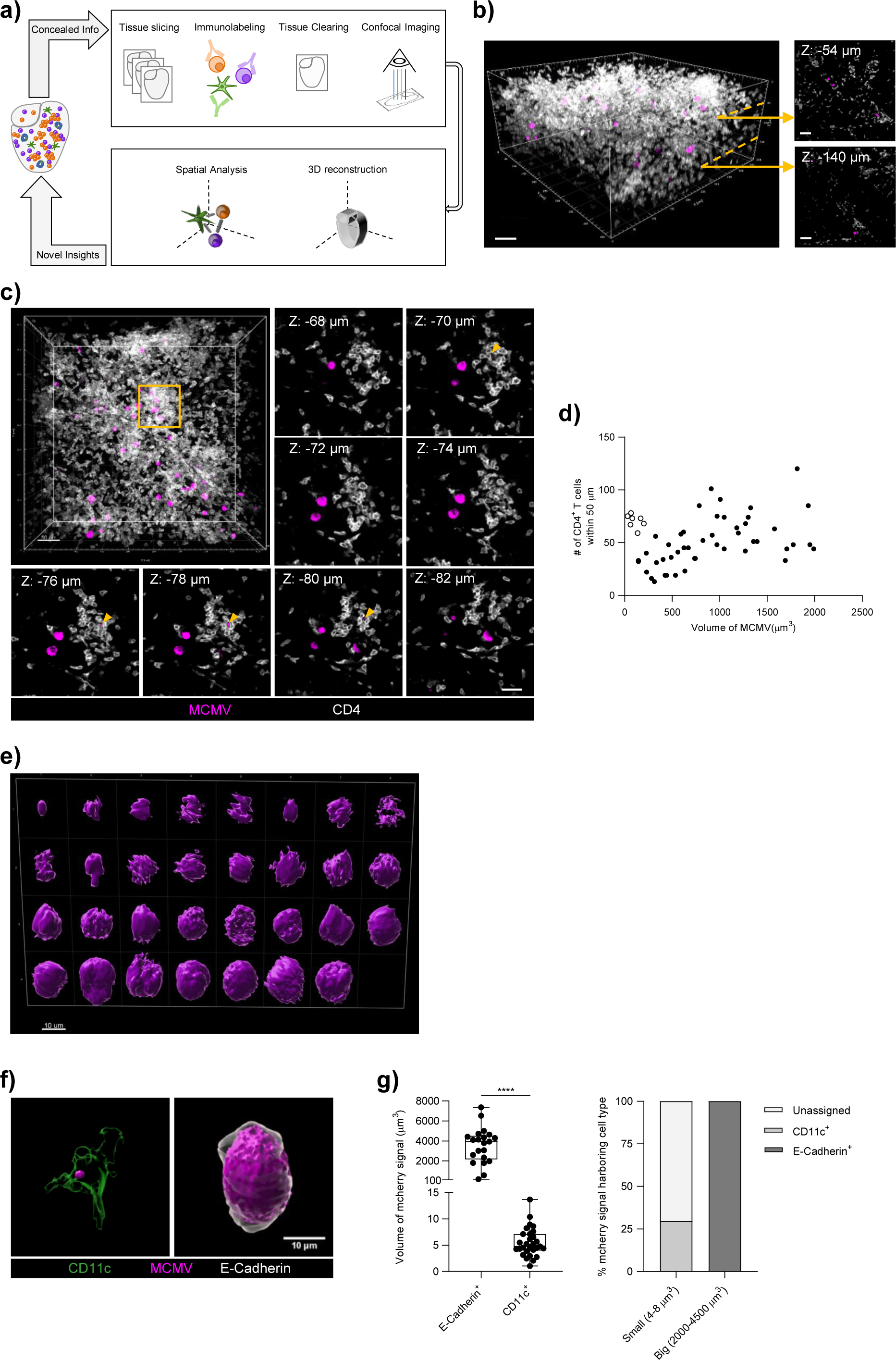
3D Imaging of MCMV-infected SGs. **a)** Schematic overview of our 3D imaging pipeline. **b)** Confocal imaging depth of an optically cleared SG sample. **c)** 3D global view (450 µm x 450 µm x 200 µm) of a 200 µm thick SG section (left, top) with magnified 2D sections at indicated z-positions from the outlined FOV (yellow frame). Yellow arrowheads show MCMV-3DR-associated faint mcherry signal. **d)** MCMV-3DR volume plotted against the number of CD4^+^ T cells within 50 µm to each mcherry signal. Dashed circle demarcates potential remnants of previously MCMV- 3DR infected cells with high abundance of CD4^+^ T cells (open dots) **e)** Morphological variability of MCMV-3DR-associated mcherry signals from FOV represented in **c**, ordered by volume from top left to down right. **f)** 3D reconstructions of MCMV-3DR-associated mcherry signal within a CD11c^+^ and an E-Cadherin^+^ cell 14 dpi. **g)** Volume of MCMV-3DR-associated mcherry signal in E-Cadherin^+^ and CD11c^+^ cells 14 dpi (left). Percentage of E-Cadherin^+^ and CD11c^+^ cells harboring small (4-8 µm^3^) and big (2000-4500 µm^3^) volumes of MCMV-3DR-associated mcherry signal 14 dpi (right). Data in **d** are shown as dots of n = 54 MCMV-3DR reporting mcherry signal pooled from 2 FOVs of one independent experiment. Data in **g** are shown as dots of MCMV-3DR reporting mcherry signals in n = 21 E-Cadherin^+^ cells and n = 28 in CD11c^+^ cells and as percentage of n = 103 small and n = 21 big MCMV-3DR reporting mcherry signals pooled from 9 FOVs of 3 independent experiments. Statistical significance was determined using single unpaired two-tailed t test **(g)**. ****P<0.0001. Scale bar in **b** = 50 µm, in **c** = 50 µm (3D global view) and 30 µm (magnified 2D FOVs), in **e** = 10 µm and in **f** = 10 µm. FOV = Field of view.

### The chemokines CXCL9 and CXCL10 act as chemoattractants in guiding CXCR3- competent T cells to site of infection

In-depth confocal imaging analysis revealed a preferred clustering behavior of CD4^+^ T cells in vicinity to sites of infection, and identified further remnants from virus-infected cells as primary antigenic source for IFN_γ_ production. To determine the range in which secreted IFN_γ_ would be sensed in this glandular tissue in order to induce an antiviral state, we analyzed the expression of the known IFN_γ_-inducible chemokines CXCL9 and CXCL10 upon MCMV infection using the CXCR3-ligand reporter mouse Rex3^30^. In Rex3 mice, CXCL9 and CXCL10 positive cells can simultaneously be detected by concomitant expression of the RFP and blue fluorescent protein (BFP), respectively. Firstly, we i.v. infected Rex3 mice with MCMV-GFP. 60% of CD45.2^+^ hematopoietic cells expressed CXCL10 in naïve SGs, which unexpectedly remained constant following viral infection (**Fig. 6a).** In contrast, only CD31^+^ blood vessel constituting endothelial cells from the CD45.2^-^ non-hematopoietic cell fraction significantly upregulated CXCL10 in response to MCMV-GFP infection, whereas CXCL10 expression levels by CD326^+^ duct forming epithelial cells and CD31^-^CD326^-^ stromal cells were hardly affected by the infection (**Fig 6a, suppl. Fig. 5a-c).** Furthermore, 70-80% of SGs-resident and half of spleen-localized CD11c^+^MHCII^+^ APCs were single CXCL10 positive, and only a minority co-expressed CXCL9 and CXCL10 (**Suppl. Fig. 5d).** Additionally, most APCs in the SGs were positive for the macrophage-specific marker F4/80 (**Suppl. Fig. 5e**). Next, we performed microscopic analysis to uncover the spatial relationship between SG-infiltrating CD4^+^ T cells, sites of viral replication, and the presence of CXCL9 and CXCL10 positive cells. Of particular importance, CD4^+^ T cells were frequently found in small discrete clusters along with CXCL9 and CXCL10-expressing cells. As observed before, these accumulations were not localized in immediate vicinity to sites of viral replication, but rather within close proximity (**Fig. 6b).** To elucidate whether CXCL9 and CXCL10 served as proxy for IFN_γ_ downstream signaling or as chemoattractants, or both, we followed two approaches. In a first experiment, we examined CXCL9 and CXCL10 expression as potential readout for IFN_γ_ sensing. For this purpose, we adoptively co-transferred either IFN_γ_-competent (IFN_γ_^+/+^) CD4^+^ and CD8^+^ T cells or IFN_γ_-deficient (IFN_γ_^-/-^) CD4^+^ and IFN_γ_-competent CD8^+^ T cells into T cell-deficient Rex3 mice (Rex3 x TCRλ3^KO^) one week prior to MCMV-GFP infection and harvested the SGs and spleens 14 or 45 dpi (**Suppl. Fig. 5f).** Fourteen days post MCMV-GFP infection, we observed similar numbers of SG-infiltrating CD4^+^ and CD8^+^ T cells (**Suppl. Fig. 5g).** Although CD4^+^ T cells account for nearly 80% of all IFN_γ_ producing cells in MCMV-infected SGs (**Fig. 4c & g)**, transfer of IFN_γ_^-/-^ CD4^+^ T cells did not significantly alter CXCL9 and CXCL10 expression levels in diverse non-hematopoietic cell types in the SGs as well as in CD11c^+^MHCII^+^ APCs in the SGs and spleens (**Suppl. Fig. 6a & b)**. Moreover, IFN_γ_^-/-^ CD4^+^ T cells still assembled together with CXCL9 and CXCL10 expressing cells, developing distinct clusters (**Fig. 6c).** The lack of IFN_γ_^+/+^ CD4^+^ T cells resulted only in a slight increase in viral loads 14 days post MCMV-GFP infection with no significant difference at 45 dpi, shortly after the peak of the infection in the SGs (**Fig. 6d, suppl. Fig. 1h**). In a second experiment, we explored the role of CXCL9 and CXCL10 as possible chemoattractants for CXCR3 expressing T cells. By using a competitive model, we investigated the infiltration and localization properties of CXCR3-competent (CXCR3^+/+^) versus CXCR3- deficient (CXCR3^-/-^) CD4^+^ T cells in the SGs upon MCMV infection. For this purpose, we adoptively co-transferred CD45.2^+^CXCR3^-/-^ and CD45.1^+^CXCR3^+/+^ CD4^+^ T cells together with CD45.1^+^CXCR3^+/+^ CD8^+^ T cells into Rex3 x TCRλ3^KO^ mice. Seven days post T cell transfer, we i.v. infected mice with MCMV and examined the SGs *in situ* at 14 dpi (**Suppl. Fig. 6c**). In line with previous results (**Fig. 6b and c**), CD4^+^ T cells were predominantly found in clearly visible accumulations, showing a very strong colocalization with CXCL9 and CXCL10 expressing cells (**Fig. 6e**). Of note, distinction between CXCR3^+/+^ and CXCR3^-/-^ CD4^+^ T cells based on the congenic marker CD45.1 demonstrated that predominantly CXCR3^+/+^CD4^+^ T cells were present within these “CXCL9 and CXCL10 hotspots”, whereas CXCR3^-/-^ CD4^+^ T cells appeared to be rather randomly distributed (**Fig. 6f**). This suggests that CXCR3 enables migration of T cells towards CXCL9 and CXCL10 positive cells within the SGs. However, CXCR3 expression was not a prerequisite for the entry into MCMV-infected SGs, and did not have an impact on the phenotype of CD4^+^ T cells (**Suppl. Fig. 6d & e).** This latter observation confirms previous findings that showed a redundant role of CXCR3 expression for T cell infiltration into MCMV-infected SGs^31^. Together, these data indicate that CXCL9 and CXCL10-expressing cells in MCMV-infected SGs possess a crucial role in guiding and positioning CXCR3 expressing T cells, and that the presence of IFN_γ_-deficient CD4^+^ T cells neither compromise expression levels of these chemokines nor have an impact on virus control around the peak of viral load in the SGs 45 dpi.

**Figure 6.**
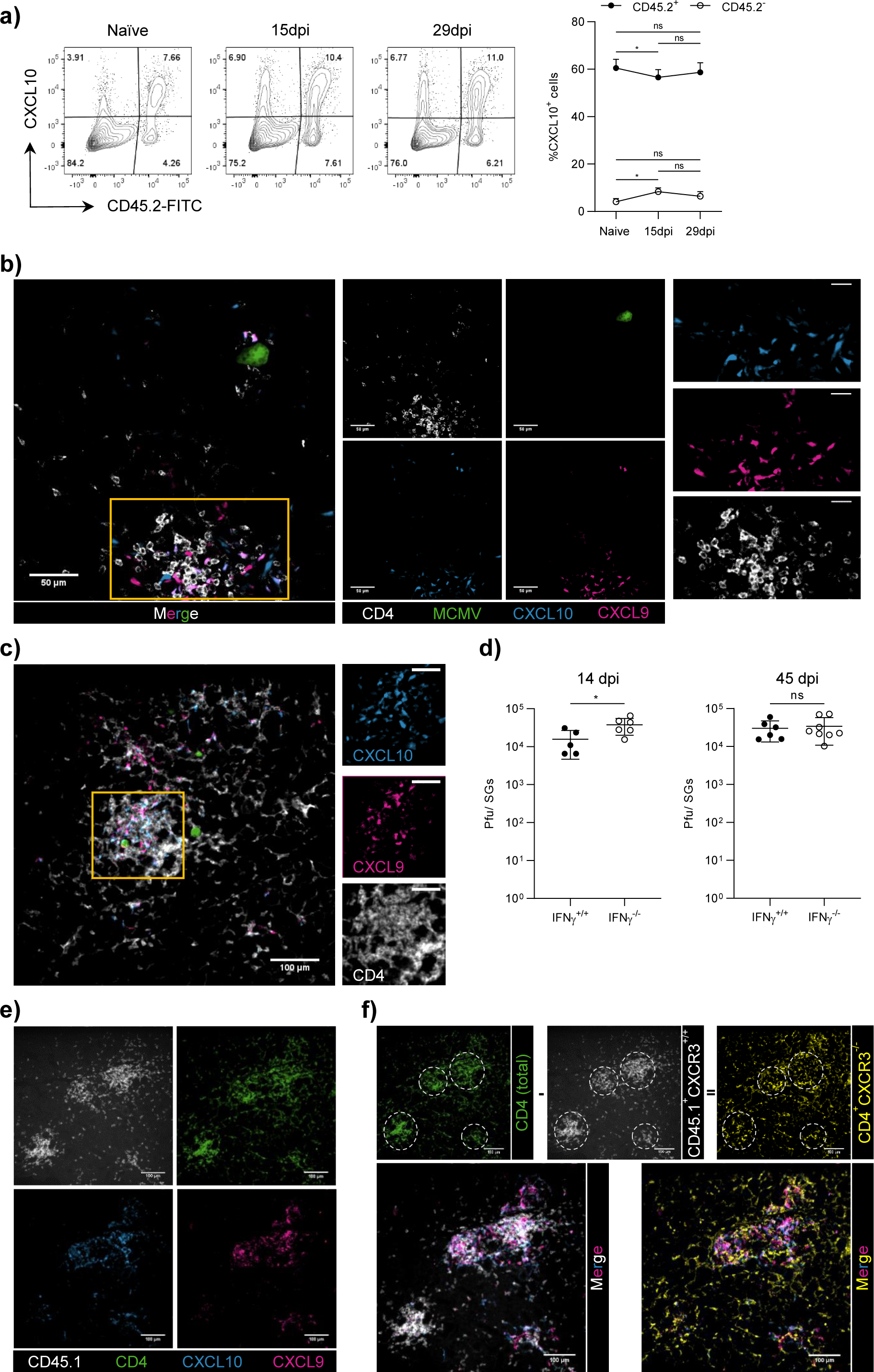
CXCL9 and CXCL10 expression in MCMV-infected SGs. **a)** Representative flow cytometry contour plots of naïve and MCMV-GFP infected Rex3 mice (left) and percentage of CXCL10^+^ hematopoietic (CD45.2^+^) and non-hematopoietic (CD45.2^-^) cells at indicated days post MCMV-GFP infection (right). **b)** Example of a four color FOV of a Rex3 mouse-derived SG 15 days post MCMV-GFP infection (left). Four single color channel images (middle) and magnified region (yellow frame) split in three single color channel images (right). **c)** Example of a four color FOV of a Rex3 x TCRλ3^-/-^ mouse-derived SG harboring IFN_γ_^-/-^ CD4^+^ T cells 14 days post MCMV- GFP infection (left) with magnified region (yellow frame) split in three single color channel images (right). **d)** Virus loads in the SGs of TCRλ3^-/-^ mice harboring IFN_γ_ competent (IFN_γ_^+/+^) or IFN_γ_ deficient (IFN_γ_^-/-^) CD4^+^ T cells 14 and 45 dpi. **e)** Example of a four single color FOV of a Rex3 x TCRλ3^-/-^ mouse-derived SG harboring CXCR3 competent (CD45.1^+^CXCR3^+/+^) and CXCR3 deficient (CD45.2^+^CXCR3^-/-^) CD4^+^ T cells. **f)** Upper row: Image subtraction. Lower row: Merged three color CD45.1^+^CXCR3^+/+^/CXCL9/CXCL10 (left) and CD4^+^CXCR3^-/-^/CXCL9/CXCL10 (right) images. Data in **a** are shown as mean + SD of n = 6 - 8 Rex3 mice pooled from two independent experiments. Data in **d** are shown as mean ± SD of n = 5 - 6 TCRλ3^-/-^ mice pooled from two independent experiments. Each dot represents either the mean of pooled mice (**a**) or one individual mouse (**d**). Statistical significance was determined using 2-way Anova with post hoc Tukey’s multiple comparisons test (**a**) or single unpaired two-tailed t test (**d**). *P<0.05, ns = not significant. Scale bar in **b** = 50 µm (left and middle) and 20 µm (right), in **c** = 100 µm (left) and 50 µm (right), in **e** and **f** = 100 µm.

### Mathematical modeling supports eventual viral control by confined local IFN*γ* production of CD4^+^ T cells in the SGs

To determine if locally confined IFN_γ_ release by CD4^+^ T cells can eventually lead to area-wide control of MCMV infection in the SGs, we combined experimental data with a mathematical model that simulates the interaction of immune and infection processes. Using a cellular Potts modeling framework that allows us to follow the dynamics and motility of individual cells over time, we recapitulated the tissue structure and main cell types within the SMG. The multi-scale model considers the turnover and infection of epithelial cells, including intracellular viral replication and extracellular viral spread, as well as the infiltration, migration and locally confined IFN_γ_ production by endogenous CD4^+^ and M25 CD4^+^ T cells (**Fig. 7a & b)**. Depending on the IFN_γ_ concentration in their surrounding, viral replication of infected cells is inhibited and infected cells are lost. Individual processes and model components were parameterized according to previous data from the literature and adapted to the experimental data. Simulating a total area of 800×800 μm^2^, our model represents roughly 1/30 of a 2D cross section of the SMG. For a detailed description of the model and its parameterization please refer to **Materials & Methods** and the detailed **Supp. Information Text S1 – Mathematical model**.

**Figure 7.**
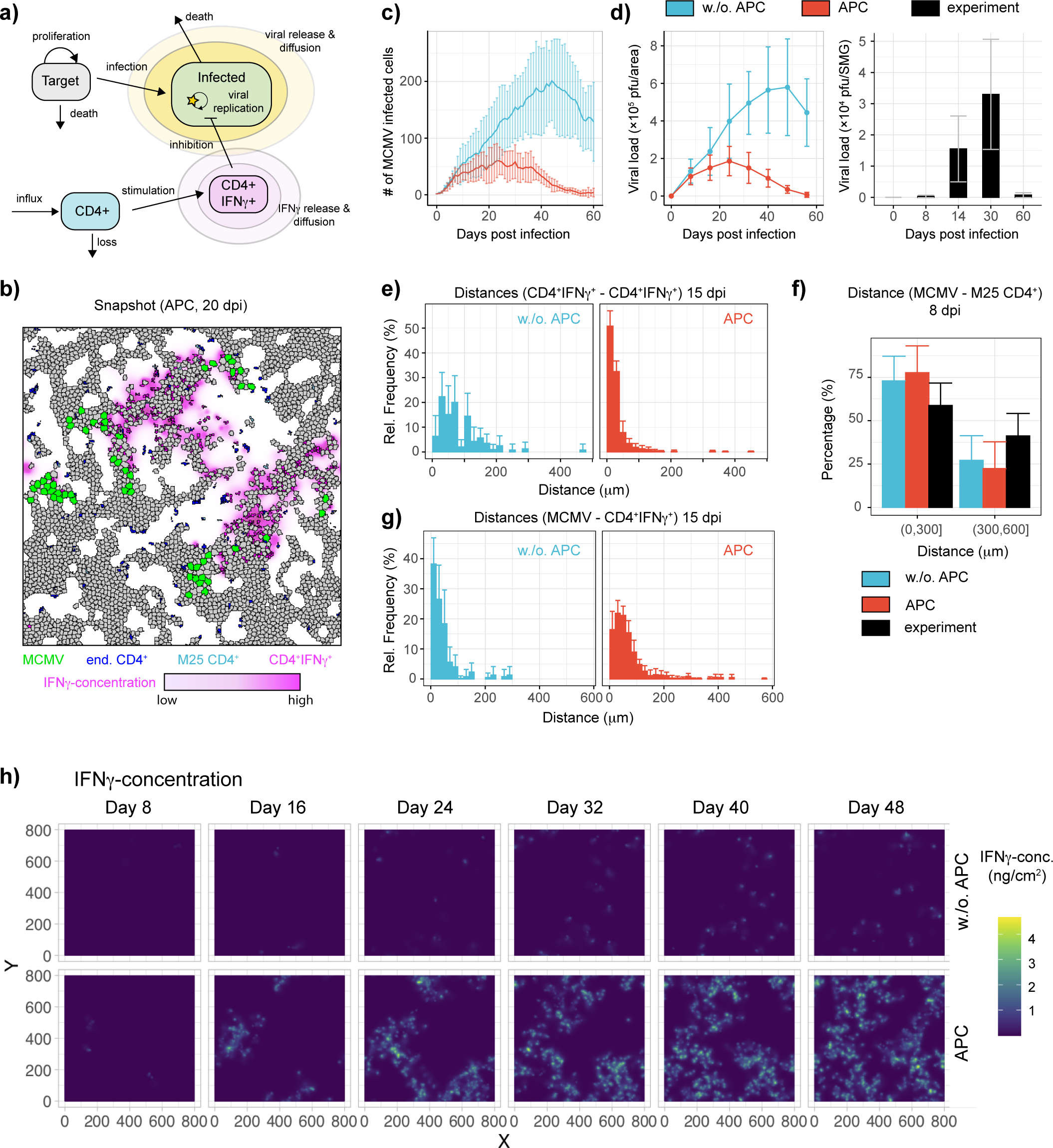
Mathematical modeling of viral-immune interactions in the SGs. **a)** Sketch of the main processes considered within the mathematical model. **b)** Snapshot of the modeled SMG tissue area of 800×800 μm^2^ at 20 dpi considering APC related guidance of CD4^+^ T cells. The plot indicates the non-duct areas with uninfected (grey) and active MCMV harboring (green) epithelial cells, as well as endogenous CD4^+^ (blue) and M25 CD4^+^ T cells (light blue). CD4^+^ IFN_γ_^+^ T cells are shown in pink with IFN_γ_^+^ concentrations indicated by the same, gradient color. **c and d)** Dynamics of the number of infected (MCMV^+^) cells **(c)** and viral load **(d)** for scenarios with (red) and without (blue) APC-associated CXCL9/CXCL10 attraction of CD4^+^ T cells. Experimental data show the viral load for the whole SMG (n= 8-9 mice, mean+/-SD). **e)** Clustering of CD4^+^ IFN_γ_^+^ T cells indicated by the distance of one IFN_γ_ producing CD4^+^ T cell to its nearest neighbor. **f)** Distances of M25 CD4^+^ T cells to the nearest MCMV infected cell. **g)** Distances of IFN_γ_ producing CD4^+^ T cells to the nearest MCMV infected cell. **h)** Dynamics of the IFN_γ_ concentration within the simulated tissue section of 800×800 μm^2^ over the progression of infection. For each scenario, i.e., w./o. and with APC, one representative simulation out of ten replicates is shown. All simulations were run over 60 days in total. Results in panel **(c-g)** are based on ten replicates for each scenario, with the mean and standard deviation (error bars) indicated. Please note that simulations are generally run with higher infection dynamics than observed experimentally for computational efficiency to account for the small simulated grid size (see Materials & Methods). Therefore, viral loads can only be qualitatively compared to experimental data.

Our simulations show that after an initial increase in the number of infected cells and total viral load, peaking around 20-30 dpi, local IFN_γ_ secretion by infiltrating CD4^+^ T cells can eventually control viral infection in the SGs 45-60 dpi (**Fig. 7c & d)**, similar to the experimental data. In this regard, a model that includes attraction of CD4^+^ T cells to CXCL9 and CXCL10 positive tissue areas populated with APCs, which engulf remnants of previously infected cells, contributes to reduced pathology and faster immune control of the MCMV infection. In contrast, a scenario that does not consider this chemical gradient-mediated migration behavior of CD4^+^ T cells, but rather assumes an undirected motion of CD4^+^ T cells (i.e. w/o APCs), is associated with an elevated and less controlled viral burden. The former scenario leads to increased clustering properties of CD4^+^ IFN_γ_^+^ T cells (**Fig. 7e**), with the majority of M25 CD4^+^ T cells accumulating in close proximity to actively MCMV infected cells (**Fig. 7f**). However, IFN_γ_-producing CD4^+^ T cells are not directly located next to sites of active viral replication, which is in line with the experimental observations (**Fig. 4e & 7g)**. Of note, IFN_γ_ producing CD4^+^ T cells do not show signs of clustering and are more heterogeneously distributed if the relevant contribution of APCs is ignored (**Fig. 7e**). Next, clustering of CD4^+^ T cells by APCs enables a concerted release of IFN_γ_ by several CD4^+^ T cells, which leads to increased local cytokine concentrations (**Fig. 7h**). In doing so, the accumulation of IFN_γ_−protected sites over time finally leads to viral control in the entire simulated tissue section.

As our model assumes a restricted effective range for IFN_γ_ released by CD4^+^ T cells with a at least 10%-chance to inhibit viral replication up to a radius of roughly ∼30 μm around the cell, individual CD4^+^ T cells are less effective in controlling active viral replication. Therefore, in case of not provoking accumulations of CD4^+^ T cells by APCs, IFN_γ_ concentrations are generally lower and more transient, mediating less efficient control of the infection (**Fig. 7c, d & h)**.

Thus, our model shows that accumulations of CD4^+^ T cells by CXCL9/CXCL10 chemotactic gradients increases the efficiency of confined IFN_γ_ release by individual CD4^+^ T cells, which confer improved regional-confined tissue protection and eventual viral control.

## Discussion

Infection of mice with MCMV induces an acute infection with lytic viral replication, followed by persistence of latent viral genomes from which sporadic reactivation events can occur. While acute infection is controlled in most organs within one to two weeks, the SGs represent a unique mucosal niche with sustained viral replication and hence persistence over several weeks^32^. Despite the marked infiltration of functional T cells, latency in this peripheral glandular tissue is only achieved two months post viral trigger. One key contribution to this delayed immune control is CMV’s interference with antigen presentation that prevents the recognition of infected cells by CD8^+^ T cells, and in less documented circumstances by CD4^+^ T cells too^33–39^. Since the decreased expression or the complete lack of MHC class I on affected cells would induce a “missing self” signal and hence render virally infected cells susceptible to NK cell-mediated killing, CMV encodes for a MHC class I homolog, serving as surrogate MHC class I molecule preventing NK cell attack^40^. Moreover, MCMV mutants lacking the m157 gene and having a functional version of MCK2 gene product, reveal enhanced dissemination and virulence properties^41–47^. Therefore, the acquisition of beneficial virus-specific *in vivo* characteristics that allow MCMV to persist preferentially in the SGs are responsible for the negligible role of NK and CD8^+^ T cells to control MCMV infection in the SGs, and further highlight a superior contribution of CD4^+^ T cells in viral control^7, 8^.

In the present study, we investigated the crucial antiviral effector functions by CD4^+^ T cells upon CMV infection in murine salivary glands. Although previous studies have already identified an indispensable role of IFN_γ_ producing CD4^+^ T cells in controlling MCMV infection in the SGs, relevant spatiotemporal information was missing that might help to explain the overall slow process of CD4^+^ T cell-mediated viral control^12, 13^.

Here, we combined experimental data together with mathematical models to provide a mechanistic understanding of the dynamics of viral spread and immune protection on a cellular level. By generating detailed micro-anatomical information about the tissue localization of infected cells, CD4^+^ T cells, sites of antigen recognition and IFN_γ_ production, and inclusion of these data into mathematical models, we propose a scenario that explains the delayed control of MCMV infection in the SGs.

We found that M25-specific CD4^+^ T cells successfully infiltrated MCMV infected SGs early upon infection, displayed an activated phenotype, and produced considerable amounts of the key pro- inflammatory cytokine IFN_γ_ upon encountering the cognate M25 antigen. However, with gradually increasing viral loads in the SGs, decreasing numbers of M25 CD4^+^ T cells, together with reduced TCR signaling events over time, suggested a negligible contribution to early MCMV control. Possible reasons might be the temporal shift from early dominant M25-specific CD4^+^ T cells to M09-specific CD4^+^ T cells at later time points^17^, and possibly a role for CD4^+^ T cells with other MCMV-specificities, given the large MCMV genome containing more than 200 open reading frames with antigenic potential^32, 48^. However, we have previously shown that the adoptive transfer of *in vivo* activated M25 CD4^+^ T cells into sublethally irradiated mice conferred immune protection in various organs in an IFN_γ_-dependent manner, indicating that M25-specific cells are able to contribute to MCMV control in immunocompromised mice^19^. Focusing on the abundant endogenous CD4^+^ T cell population within the MCMV infected SGs, we demonstrated that CD4^+^ T cells are the primary cellular source of IFN_γ_ production, in accordance to elevated TCR downstream signaling, indicative of antigen recognition. Moreover, *in situ* analysis of MCMV infected SMGs revealed that IFN_γ_ expressing CD4^+^ T cells were mostly found in vicinity to sites of lytic viral replication, in dense clusters surrounding CD11c^+^ cells that contained small inclusions of cellular cargo from previously infected cells. Clearly, these CD11c^+^ cells were not actively infected with MCMV, as this would have resulted in presence of the reporter fluorochrome within the entire cytoplasm and not within confined small areas as observed. We thus speculate that this cargo represents apoptotic cell bodies from previously infected cells^11^. In this regard, Stahl et al. made a similar observation in infected neonatal lungs, where infection with MCMV attracted myeloid cells, leading to the generation of morphologically unique “nodular inflammatory foci (NIF)” (multiple juxta-positioned MCMV infected cells and associated immune cell infiltrates). Within NIF, which were required for efficient containment of the infection, they characterized few small mcherry-containing apoptotic bodies within myeloid cells, revealing the possible uptake of cell debris derived from previously infected cells by these cells^49^. As we observed discrete accumulations of CD4^+^ T cells around CD11c^+^ APCs, with some of them revealing cargo of MCMV infected cells, in combination with local TCR triggering and IFN_γ_ production, we were wondering which signals are required for the recruitment of these cells into regions of antigen presentation. We specifically investigated the role of the CXC chemokines CXCL9 (MIG) and CXCL10 (IP-10) in this process. These two chemokine ligands share the common receptor CXCR3, which is predominantly expressed on effector T cells^22, 23^. Whereas IFN_γ_ signaling strongly induces CXCL9 expression, CXCL10 can also be upregulated by type I interferons (IFNα/λ3)^50, 51^. CXCR3 is important for directional migration of CD4^+^ and CD8^+^ T cells in various infectious settings^52–54^. However, a clear role of CXCR3 expression in MCMV-specific T cells has so far not been demonstrated. Although this chemokine receptor promoted the recruitment of antigen-specific CD8^+^ T cells to MCMV infected liver^55^, the lack of CXCR3 expression on CD8^+^ T cells did not interfere with their ability to infiltrate into the SGs upon MCMV infection^31^. In our study, we confirmed that CXCR3 expression on CD4^+^ T cells is dispensable for SG infiltration, however, it played a decisive role for the spatial localization within MCMV infected SGs. We show that CXCR3-competent CD4^+^ T cells are specifically recruited to sites of CXCL9 and CXCL10 expression, leading to the formation of distinct CD4^+^ T cell clusters surrounding APCs. In contrast, despite similar infiltration capacity and phenotypic characteristics of CXCR3-deficient CD4^+^ T cells, these cells rather distributed randomly. Since we identified TCR engagement and IFN_γ_ production mainly within these dense aggregations of T cells, this observation suggests that CXCR3-deficient CD4^+^ T cells might have reduced chances for antigen encounter and hence ability to produce IFN_γ_. Interestingly, surface expression of CXCR3 was almost completely absent on SG-residing CD4^+^ T cells, most likely due to ligand engagement and receptor internalization, as shown in a recent publication^56^.

Finally, by feeding our experimental data into mathematical models, we developed a simulation scenario of the CD4^+^ T cell-mediated MCMV control in the SGs. In doing so, we could show that a local rather than long-range protection capability of IFN_γ_ producing CD4^+^ T cells leads to eventual infection control in the SGs. Individual small areas, defined by IFN_γ_−producing and sensing cells, provide site-specific antiviral protection, and allow MCMV to spread and replicate at spots that have not yet been yet exposed to IFN_γ_. Furthermore, due to the indirect antigen recognition on CD11c^+^ APCs that have engulfed cargo of previously infected cells, effector CD4^+^ T cell responses occur in a delayed manner, i.e. only after a virus-infected cell has already ceased to produce new virions. This indirect antigen recognition enables ongoing viral replication over several weeks in the SGs. However, the accumulation of effector CD4^+^ T cell responses by APCs potentiates the IFN_γ_ concentration within local regions, leading to larger areas of IFN_γ_− mediated protection and faster viral control in comparison to scenarios without APC mediated clustering of CD4^+^ T cells. Ultimate control of the lytic infection and the transition into a latent state would hence only happen after continued accumulation of locally protected areas, finally resulting in an organ-wide protection. Pursuing the idea of a short-range diffusion property of IFN_γ_, Müller et al. revealed a few years ago a confined action radius of IFN_γ_ that induced potent protection mechanisms beyond the immunological synapse, in (bystander) cells located more than 80 µm away from the T cell-APC interaction^57^. These would imply that within the vicinity of IFN_γ_ producing CD4^+^ T cells, depending on the number of IFN_γ_ producing CD4^+^ T cells and the amount of per- cell produced IFN_γ_, surrounding cells being maximally eight cell layers away from the IFN_γ_ source should be able to sense the secreted IFN_γ_ and thus being protected from MCMV infection.

In this context, our mathematical model provides first mechanistic insights into the dynamics of MCMV infection and clearance in the SGs by combining different types of experimental data and measurements. Although the model only captures a small fraction of the SMG, and therefore considers a higher infection level than observed experimentally, it provides a qualitative representation of the observed infection dynamics, as well as quantitative analysis of the spatiotemporal relationships of individual cells. With the results shown above relying on parameterizations that favor local viral spread^58^, eventual viral control is also achieved by locally mediated IFN_γ_ protection if MCMV would spread more heterogeneously within the tissue given higher viral diffusion, but would require larger local IFN_γ_ perimeters. In general, the ability and rapidness of viral control depends on the extension and duration of local IFN_γ_ concentrations, which is considerably enhanced by APC-mediated clustering of CD4^+^ T cells. More detailed and time-resolved measurements, as e.g. on the stability and longevity of APC-formed CD4^+^ T cell clusters, or on the dynamics of IFN_γ_ secretion by individual CD4^+^ T cells, will help to improve the parameterization of the model and validate previous parameter choices.

## Materials & Methods

### Ethics Statement

This study was conducted in accordance to the guidelines of the animal experimentation law (SR 455.163; TVV) of the Swiss Federal Government. The protocols were approved by the Cantonal Veterinary Office of the canton Zürich, Switzerland (animal experimental permissions: 146/2014 and 114/2017).

### Mice

Eight to twelve-weeks old age -and sex-matched mice were used for each experiment described in this study. C57BL/6J mice were purchased from Janiver Elevage and referred to as WT B6 mice. C57BL/6N-Tg(TCRaM25,TCRbM25)424 Biat (M25-III) (M25xLy5.1) mice harbor TCR tg CD4^+^ T cells specific for the M25411-425 peptide on the CD45.1 congenic background. The congenic CD45.1 WT B6 mice were bred in house. Nur77GFP (JAX stock # 016617), DssRed.T3 (JAX stock # 006051), Great (JAX stock # 017580), IFNγ^-/-^ (JAX stock # 002287) and TCRβ^-/-^ (JAX stock # 002118) mice were obtained from the Jackson Laboratory. CXCR3^-/-^ (JAX stock # 005796) mice were obtained from the Swiss immunological mouse repository (SwimmR). M25xNur77GFP and M25xRFP CD4^+^ T cells were generated by crossing the M25xLy5.1 mice to the Nur77GFP -and DssRed.T3 mice, respectively. Rex3 mice were kindly provided by Prof. Matteo Iannacone (IRCCS San Raffaele Scientific Institute, Milan, Italy) in agreement with Prof. Andrew D. Luster (Massachusetts General Hospital, Massachusetts, USA), and crossed to TCRβ^-/-^ mice for the development of the Rex3 x TCRβ^-/-^ mice. CD11cYFP-Prox1mOrange2 mice were kindly provided by Prof. Cornelia Halin Winter (Swiss Federal Institute of Technology Zurich, Zurich, Switzerland). All mice were bred and housed under specific pathogen-free conditions in animal facilities at the Swiss Federal Institute of Technology in Zurich, Hönggerberg. Mice were exposed to a 12:12 h light-dark cycle with unrestricted access to water and food.

### Viruses and infections

All recombinant MCMVs were deficient in m157 protein expression (Δm157) and had a functional version of the MCK2 gene product (MCK2^+^). MCMV-GFP expresses green fluorescent protein (GFP) under the m157 promoter and MCMV-3DR expresses the red fluorescent protein mcherry under the major immediate-early promoter^59–61^. MCMV-3DR was obtained from Prof. Martin Messerle (Hannover Medical School, Hannover, Germany) in agreement with Prof. Reinhold Förster and Dr. Stephan Halle (Hannover Medical School, Hannover, Germany). The non-fluorescence encoding MCMV strain was kindly provided by Prof. Luka Cicin-Sain (Helmholtz Centre for Infection Research, Braunschweig, Germany) in agreement with Prof. <u>Stipan Jonjić</u> (University of Rijeka, Rijeka, Croatia) and referred to as MCMV^62^. MCMV strains were propagated on M2-10B4 cells and subsequently purified by ultracentrifugation on a 15% sucrose cushion as previously described^3^. Viral titers were determined by standard plaque assay on M2-104 cells with centrifugal enhancement. Mice were infected with 2×10^5^ or 5×10^5^ plaque forming units (Pfu) of the respective MCMV mutant by i.v. tail injection.

### Adoptive transfers

CD4^+^ T cells from naïve M25xLy5.1, M25xNur77GFP, M25xRFP, IFNγ^-/-^, CXCR3^-/-^ or CD45.1 WT B6 mice, and CD8^+^ T cells from naïve CD45.1 or CD45.2 WT B6 mice, were purified from spleens by positive selection using anti-CD4 or anti-CD8α magnetic activated cell sorting (MACS) beads (Miltenyi Biotech, Bergisch Gladbach, Germany) according to the manufacturer’s instructions. Cell viability was assessed by trypan blue exclusion and confirmed to be greater than 90%. Flow cytometric cell analyses indicated >95% purity of the isolation procedures. 9×10^5^ or 10^6^ M25 CD4^+^ T cells were adoptively transferred into naïve recipient WT B6 mice one day prior MCMV or MCMV-GFP infection. 3×10^6^ or 3.5×10^6^ CD4^+^, and 10^6^ CD8^+^ T cells were adoptively transferred into naïve recipient Rex3 x TCRβ^-/-^ or TCRβ^-/-^ mice seven days prior MCMV or MCMV-GFP infection. Cell transfers were executed by i.v. tail injection.

### *In vivo* peptide stimulations & intravascular antibody labeling

10^6^ MACS isolated M25 CD4^+^ or M25xNur77GFP CD4^+^ T cells were adoptively transferred one day prior MCMV infection into naïve recipient WT B6 mice. M25 peptide (amino acid sequence: NHLYETPISATAMVI) was purchased from EMC microcollections (Tübingen, Germany) and 20 μg of the M25 peptide in 0.5% DMSO in 1x PBS solution were injected i.v. at day 8 or 21 post MCMV infection. Three hours post M25 peptide challenge, SGs and spleens were harvested in complete cell culture medium (RPMI1640 supplemented with 10% FCS, 2mM L-Glutamine, 1x penicillin/streptomcyin) with additional 10 μg/ml Brefeldin A. Following digestion or meshing of organs was performed in further presence of 5 μg/ml Brefeldin A.

Intravascular labeling was performed by i.v. administration of 3 μg of fluorophore-conjugated anti- CD45.1 antibody 3 min prior euthanasia with CO2 as described previously^63^. Organs were subsequently perfused with 10 ml of 1x PBS and harvested for subsequent lymphocyte isolation and flow cytometric analysis.

### Lymphocyte isolation and Flow cytometry

Mice were euthanized by CO2 inhalation, and the SGs and lungs were subsequently perfused via the right heart ventricle with 10 ml ice-cold 1x PBS to remove circulating blood. Single-cell suspensions of spleens and lungs were prepared as previously described with slight modifications^64^. Briefly, spleens were passed through metal grids using syringe plungers. The SGs and lungs were cut into small pieces and digested in complete cell culture medium containing 2.4 mg/ml collagenase type I and 0.2 mg/ml DNaseI for 2 x 20 min at 37°C on a thermomixer. Additionally, they were flushed through 18G and 21G needles in between and after the second round of digestion, respectively, and subjected to centrifugation over a 30% Percoll gradient.

Prior to fluorophore-conjugated antibody labeling, cells were treated with a hypotonic ammonium-chloride-potassium lysis buffer for 5 - 10 min at room temperature (RT) to remove remaining erythrocytes. Cells were washed and cell surface staining was executed for 20 min at RT in the dark. In case of continuing intracellular cytokine staining, cells were fixed and permeabilized in 2 × FACS BD Lysis Solution (BD Biosciences) with 0.08% Tween20 in ddH2O for 10 min at RT in the dark. Subsequent intracellular staining for IFNy and TNFα was performed for 35 min at RT in the dark. Dead cells were excluded by applying a LIVE/DEAD fixable Near-IR staining (ThermoFisher Scientific). All staining and washing steps were performed in FACS buffer (2% FCS and 5mM EDTA in 1x PBS). Fluorescently labeled antibodies were purchased from Biolegend (Lucerna Chem AG) or BD Biosciences, and are summarized in **Table 1**. Multi-parametric flow cytometric analysis was performed using LSRII flow cytometer (BD Biosciences, Allschwil, Switzerland) with FACSDiva software. Data were analyzed using FlowJo software (Version 10.4.2, Tree Star).

**Table 1.** Flow cytometry antibody list (Surface & **Intracellular staining**)

| Marker | Fluorochrome | Manufacturer | Clone | Catalog # | RRID | Conc |
| --- | --- | --- | --- | --- | --- | --- |
| B220 | PE-Cy7 | Biolegend | RA3-6B2 | 103222 | AB_313005 | 0.4 µg/ml |
| B220 | BV510 | Biolegend | RA3-6B2 | 103247 | AB_2561394 | 1 µg/ml |
| B220 | FITC | Biolegend | RA3-6B2 | 103206 | AB_312991 | 1 µg/ml |
| B220 | PE | Biolegend | RA3-6B2 | 103208 | AB_312993 | 0.4 µg/ml |
| B220 | APC | Biolegend | RA3-6B2 | 103212 | AB_312997 | 0.4 µg/ml |
| CD11a/CD18<br>(LFA-1) | PE-Cy7 | Biolegend | H155-78 | 141011 | AB_2564307 | 1 µg/ml |
| CD11c | FITC | Biolegend | N418 | 117306 | AB_313775 | 2.5 µg/ml |
| CD279<br>(PD-1) | PE-Cy7 | Biolegend | 29F.1A12 | 135215 | AB_10696422 | 1 µg/ml |
| CD3 | PE-Cy7 | Biolegend | 145-2C11 | 100320 | Ab_312685 | 1 µg/ml |
| CD3 | FITC | Biolegend | 145-2C11 | 100306 | AB_312671 | 1 µg/ml |
| CD31 | PE-Cy7 | Biolegend | 390 | 102418 | AB_830757 | 0.3 µg/ml |
| CD326<br>(EpCAM) | AF647 | Biolegend | G8.8 | 118212 | AB_1134101 | 0.8 µg/ml |
| CD4 | BV421 | Biolegend | RM4-5 | 100544 | AB_11219790 | 1 µg/ml |
| CD4 | BV605 | Biolegend | RM4-5 | 100548 | AB_2563054 | 1 µg/ml |
| CD44 | BV510 | Biolegend | IM7 | 103044 | AB_2650923 | 1 µg/ml |
| CD45.1 | PE | Biolegend | A20 | 110708 | AB_313497 | 0.4 µg/ml |
| CD45.1 | APC | Biolegend | A20 | 110714 | AB_313503 | 0.4 µg/ml |
| CD45.2 | FITC | Biolegend | 104 | 109806 | AB_313443 | 0.4 µg/ml |
| CD49d | FITC | Biolegend | R1-2 | 103606 | AB_313037 | 2.5 µg/ml |
| CD62L | PerCP | Biolegend | MEL-14 | 104430 | AB_2187124 | 1 µg/ml |
| CD69 | PE-Cy7 | Biolegend | H1.2F3 | 104512 | AB_493564 | 1 µg/ml |
| CD69 | FITC | BD<br>Biosciences | H1.2F3 | 55326 | . | 2.5 µg/ml |
| CD8a | PerCP | Biolegend | 53-6.7 | 100732 | AB_893423 | 1 µg/ml |
| CD8a | APC | Biolegend | 53-6.7 | 100712 | AB_312751 | 1 µg/ml |
| CXCR3 | BV421 | Biolegend | CXCR3-173 | 126521 | AB_10900974 | 1 µg/ml |
| F4/80 | AF647 | Biolegend | BM8 | 123122 | AB_893480 | 2.5 µg/ml |
| I-A/I-E<br>(MHCII) | PerCP | Biolegend | M5/114.15.2 | 107624 | AB_2191073 | 0.4 µg/ml |
| NK1.1 | PE | Biolegend | PK136 | 108708 | AB_313395 | 1 µg/ml |
| NK1.1 | PE-Cy7 | Biolegend | PK136 | 108714 | AB_389364 | 1 µg/ml |
| **IFNγ** | **APC** | **Biolegend** | **XMG1.2** | **505810** | **AB_315404** | **1 µg/ml** |
| **TNFα** | **FITC** | **Biolegend** | **MP6-XT22** | **506304** | **AB_315425** | **5 µg/ml** |

### Plaque forming assay

Viral loads in organs were determined on M2-104 cells as described previously^65^. Briefly, 50’000 M2-10B4 were added to each well of a 24-well plate one day prior organ-derived virus exposure. Freshly harvested or in -80°C stored organ in 1 ml complete cell culture medium was homogenized using a metal bead at 25 Hz for 1 - 1.5 min by TissueLyser (Qiagen, Hilden, Germany). After centrifugation at 3500g for 5 min, virus containing supernatant was taken and transferred to a new Eppendorf tube. A 10x serial dilution of the supernatant in complete cell culture medium was performed using round-bottom 96-well plates. Medium of the previous day cultivated M2-10B4 cells was aspirated and replaced by the diluted supernatant. The 24-well plates were centrifuged for 30 min at 535g and subsequently incubated for 1h at 37°C. Medium was aspirated and replaced by 2.2 % Methylcellulose (4000 centipoise) in ddH2O supplemented with 1x MEM, 10% FCS, 2mM L-Glutamine, 1% penicillin-streptomcyin and 10mM HEPES/sodium bicarbonate solution. Plates were incubated for four days at 37°C. After four days of incubation, 500 μl of 25% Formaldehyde solution in ddH2O was added on top of the methylcellulose layer and plates were left for 3h at RT. Next, plates were inverted to remove viscous medium and carefully washed three to four times with 1x PBS. Crystal violet solution was added to the wells and plates were incubated for 20 min at RT. Finally, plates were washed with ddH2O, air-dried, and plaques were counted using an inverted microscope. With the exception of the 1h incubation period at 37°C, all steps were carried out at 4°C. Detection limit was set to 100 Pfu.

### 2D Immunohistochemistry - Tissue preparation & Staining

1x PBS-perfused SGs were fixed in methanol-free 4% formaldehyde solution for 4 - 6 h at 4°C. After a short wash in 1x PBS, organs were cryoprotected in 30% sucrose in 1x PBS overnight (o/n) at 4°C, followed by an embedding in optimal cutting temperature (O.C.T) compound and a snap-freezing in liquid nitrogen. The SG samples were stored at -80°C until further usage. Frozen tissue sections of 10 µm thickness were generated by a cryostat-microtome, 2 - 3 hours air-dried, and stored at -20°C if not directly used. Completely air-dried thin tissue sections of the SGs were washed with 1x PBS to remove remaining O.C.T compound and marked with an hydrophobic barrier PAP pen (H-4000, Vector Laboratories). Tissue sections were blocked in 1x PBS containing 10% normal goat serum (NGS), 1% Bovine serum albumin (BSA) and 0.1% TritonX- 100, for 1h at RT. Subsequent staining steps with unlabeled and/or fluorochrome-conjugated antibodies were executed for 1h at RT or o/n at 4°C in the dark. Slides were washed in between three times for 7 min in 1x PBS. Finally, sections were washed again in 1x PBS and carefully rinsed with ddH2O, before counterstained with 4’,6’-diamidino-2-phenylindole (DAPI) for 5 min at RT and mounted in home-made Mowiol-DABCO solution (see CSH protocols; RI: 1.45 - 1.52). Slides were stored at RT for one to two days in the dark before microscopic investigation. Immunolabeling was performed in 1x PBS containing 3% BSA and 0.1% TritonX-100 in a humidity chamber. Antibodies were purchased from Biolegend (Lucerna Chem AG), Life Technologies (Thermo Fisher Scientific), BD Biosciences, Clontech Laboratories and Rockland Immunochemicals, and are summarized in **Table 2**.

**Table 2.**
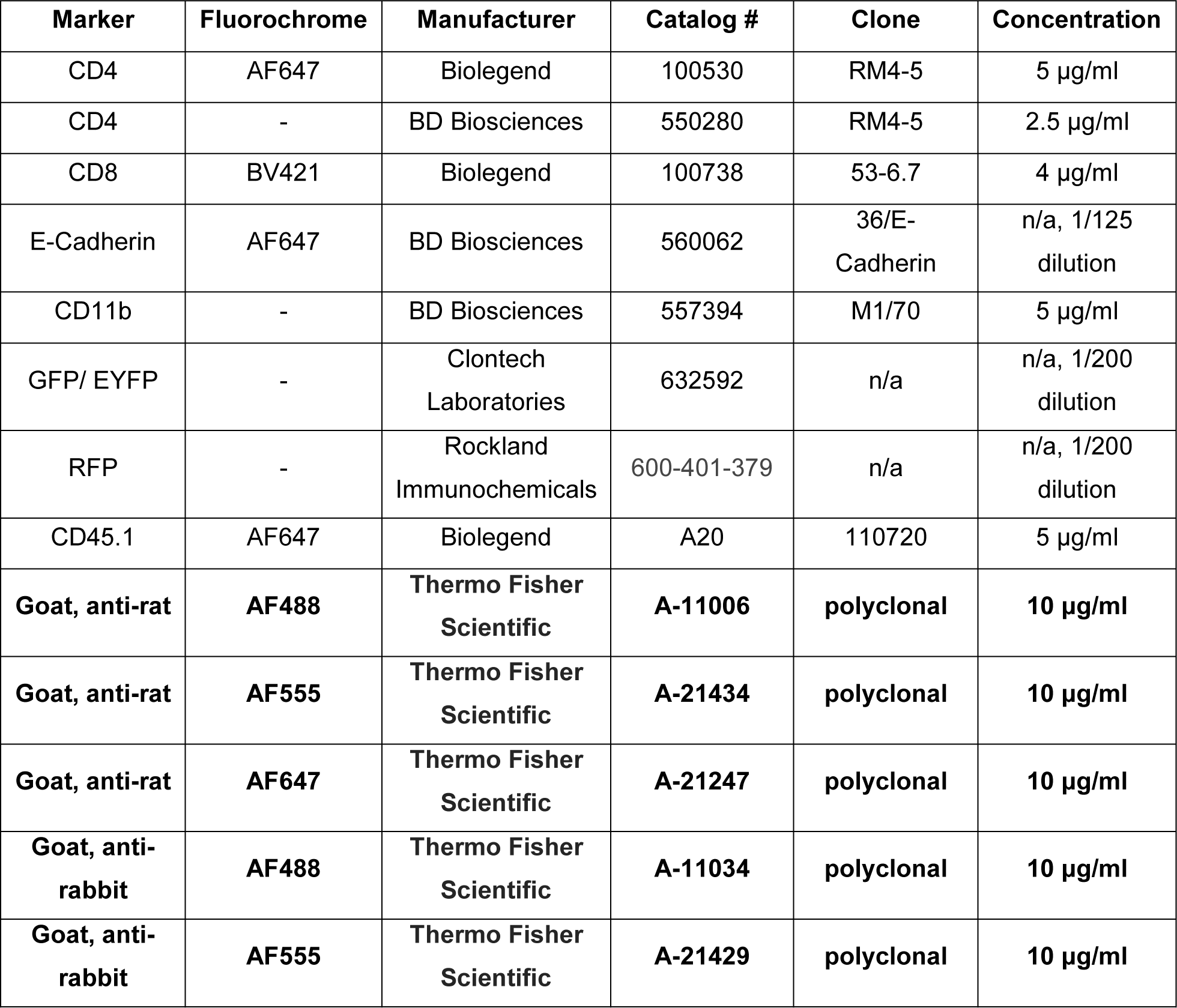
Microscopy antibody list (primary & **secondary**)

### 2D Immunohistochemistry - Imaging & Analysis

Images of thin tissue slices were acquired with an inverted confocal microscope (Axiovert 200m microscope, Carl Zeiss, Inc., Zurich, Switzerland), equipped with an CSU-X1 spinning-disk confocal unit (Yokogawa), a 20x (Nummerical aperture (NA): 0.5, Immersion: Air) or a 10x (NA: 0.3, Immersion: Air) magnification phase contrast objective, a solid state laser unit with 4 laser lines (405, 488, 561, 647, Toptica) and two Evolve 512 EMCCD cameras (Photometrics). Detection and spectral separation of excited fluorophores were achieved through the usage of specific emission bandpass filters (450/50, 525/50, 605/52 and 700/75). Images with sequential fluorophore excitation and with z-stacks sizes of 1 µm were recorded and analyzed using imageJ/Fiji software (Version 1.51). An initial brightness/contrast enhancement was performed with subesequent quantitative analysis of IFN_γ_ producing (EYFP^+^) cells using the cell counter plugin.

### Whole slide Imaging & Analysis

10 µm thin frozen tissue sections of MCMV infected SGs were prepared and stained according to the 2D immunohistochemistry part. However, only SG sections, which do not show any signs of processing-mediated artifacts (e.g. tissue distortions, air bubbles or folds), were subjected to whole slide scanning using Slide Scanner Pannoramic 250 Flash III (3D Histech, Budapest, Hungary) to realize faithful data for further downstream analysis. Images with a z-stack size of 0.6 µm were acquired using a 20x magnification objective (NA: 0.8, Immersion: Air). Recorded mrxs files were directly imported into the HALO^®^ software (https://indicalab.com/halo/; Version 3.0.311.373; Indica Labs, Albuquerque, New Mexico, USA) for quantitative image analysis. In doing so, we first applied tissue annotation to delimit our spatial analysis to the SMG. We then performed a categorization of diverse tissue types (i.e. endogenous CD4^+^ and M25 CD4^+^ T cells, sites of viral replication, duct cells, other cells and background) using tissue classifier (Random Forest). We next exerted Highlex FL module (Version 3.1.0.0) for the simultaneous detection of various cell types based on fluorescence marker intensities in the different cellular compartments (i.e. nucleus, cytoplasm and cell membrane). In this regard, we first identified and segmented all DAPI^+^ cells, irrespective of cell type (Nuclear contrast threshold = 0.5, Minimum Nuclear Intensity = 0.1, Nuclear Segmentation Aggressiveness = 0.8 and Minimum Nuclear Roundness = 0). Afterwards, endogenous CD4^+^ and M25 CD4^+^ T cells were defined within DAPI^+^ cells based on fluorescence signal intensities in the previously mentioned cellular compartments (see table below). In this regard, endogenous CD4^+^ T cells expressed only cell membrane marker CD4 (= DAPI^+^CD4^+^ cells) whereas M25 CD4^+^ T cells were additionally positive for the cytoplasmic RFP signal (= DAPI^+^CD4^+^RFP^+^ cells). Finally, quantitative spatial tissue analysis was executed using nearest neighbor analysis in conjunction with proximity analysis to determine the relative spatial distribution of cells and sites of viral replication across the tissue section. Micro-anatomical localization of sites of viral replication was provided by the tissue classifier.

Tresholding (signal intensities):

| Fluorescent label<br>(marker) | Nucleus | Cytoplasm | Cell membrane |
| --- | --- | --- | --- |
| DAPI | 1 | -1 | -1 |
| AF568 (RFP) | 40 | 40 | 0 |
| AF647 (CD4) | 35 | 35 | 0 |

Signal range: 0 (= no signal) - 255 (= highest intensity)

### 3D Immunohistochemistry - Tissue preparation & Staining

1x PBS perfused SGs were fixed in methanol-free 4% formaldehyde solution for 4 - 6 h at 4°C. After a short wash in 1x PBS, organs were embedded in low-melting 4% agarose in 1x PBS, and subsequently cut in 200 µm thick tissue sections using a Compresstome^®^ (Precisionary, Greenville, USA). Slices were collected in 6-well plates pre-filled with 0.1% PBST and incubated for 24 h at RT. PBST was aspirated and sections were incubated overnight in blocking buffer (1x PBS containing 10% NGS, 1% BSA and 0.1% TritonX-100) at 4°C. Blocking buffer was removed and sections were stained in 6, 12 or 24-well plates with primary fluorochrome-conjugated antibodies diluted in 3% BSA in 1x PBS for three to four days at 4°C or RT (**Table 2).** Next, slices were washed three to four times with 1x PBS during a 24 h period at 4°C. 1x PBS was removed, and a home-made RIMS with RI = 1.47 (45 g Histodenz in 30 ml 1x PBS) was added to the sections. The samples were incubated in the dark on a rotator for 6 h at RT. Finally, SG sections were mounted using fresh RIMS and appropriate 200 µm thick iSpacer^®^ (SunJin Lab, Hsinchu, Taiwan). Tissue samples were stored one to two days at RT in the dark prior confocal imaging.

### 3D Immunohistochemistry - Imaging & Analysis

Images of 200 µm thick tissue sections were acquired with an inverted confocal microscope (Leica TCS SP8 MP), using a 25x magnification objective (NA: 0.95, Immersion: Water), a laser unit with AOBS system (405, 488 (Argon Laser), 561, 633), and internal detection with 1 PMT detector and 2 Leica Hybrid detectors, operated by the Leica LAS X 3.5.5. Images were acquired with sequential fluorophore excitation, z-stacks sizes of 2 µm and a scan format of 512x512 pixels.

Image analysis was performed using imageJ/Fiji software (Version 1.51), Imaris 9.5 (Bitplane, Zurich, Switzerland) and RStudio (Version 1.1.463). Using Imaris, several processing steps were applied to the images (**Table 3**). The acquired signals were attributed to each individual cell (surface creation), or its location was determined without accurate segmentation of cell bodies by the generation of spherical objects centered on any pixels with signal above a user set threshold (spots creation) (**suppl. Fig 4c**). For the CD4 channel, gamma was adjusted to 1.35, and Gaussian smoothing was used with a width of 0.32 µm. Finally, surfaces and spots were created according to the parameters in **Table 4**. Appeared extra-cellular and likely represented imaging artefacts were not selectively removed from analysis to avoid introducing user bias.

**Table 3.**
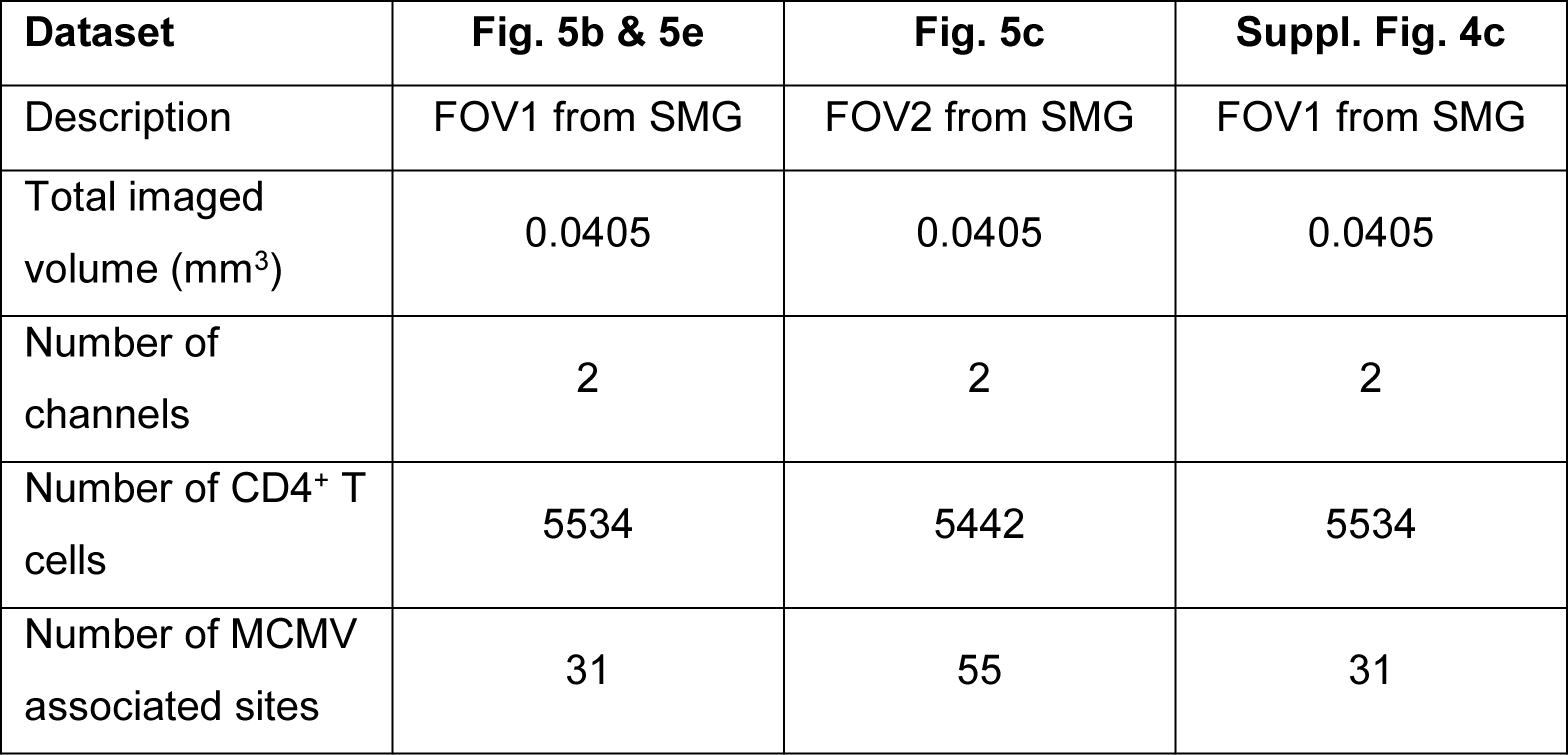
General information to volumetric images

| Dataset | Fig. 5b & 5e | Fig. 5c | Suppl. Fig. 4c |
| --- | --- | --- | --- |
| Description | FOV1 from SMG | FOV2 from SMG | FOV1 from SMG |
| Total imaged volume (mm<sup>3</sup>) | 0.0405 | 0.0405 | 0.0405 |
| Number of channels | 2 | 2 | 2 |
| Number of CD4<sup>+</sup> T cells | 5534 | 5442 | 5534 |
| Number of MCMV associated sites | 31 | 55 | 31 |

**Table 4.** Technical information to volumetric images

| Surface/ Spots creation parameters |  |  |  |  |  |  |  |  |
| --- | --- | --- | --- | --- | --- | --- | --- | --- |
| Surface/<br>Spots<br>name | Source<br>Channel | Smoothing:<br>Surface<br>details (µm) | Background<br>Subtraction:<br>Diameter of<br>largest sphere | Absolute<br>intensity<br>threshold | Split touching<br>objects: Split<br>seed diameter<br>(µm) | Quality<br>threshold | Voxel<br>number<br>threshold | Diamers<br>(spots) |
| Suppl. Fig. 7a |  |  |  |  |  |  |  |  |
| MCMV<br>surfaces | MCMV-3DR | 0.3 | 9.00 | 11.6 | 7 | 3.29 | 19-1160 | - |
| CD11c<br>surfaces | CD11c<br>eYFP | 0.5 | 10 | 27.9 | 10 | 6.03 | 1075-10060 | - |
| CD4<br>spots | CD4<br>AF647 | - | true | - | - | 2 | - | 5.5 |

### *Ex vivo* live imaging

Mice were euthanized by CO2 inhalation, and the SGs were harvested in complete cell culture medium (RPMI1640 supplemented with 10% FCS, 2mM L-Glutamine, 1x penicillin/streptomycin). The organs were embedded in 4% agarose in 1x PBS, and subsequently cut in 200 µm thick tissue sections using a Compresstome^®^ (Precisionary, Greenville, USA). Slices were collected in 6-well plates filled with live imaging medium (RPMI 1640 without phenol red supplemented with 10% FCS, 2mM L-Glutamine, 1x penicillin/streptomycin, 1M Hepes, 50 mM *β*-Mercaptoethanol, 100x Sodium Pyruvate, 100x Minimum Essential Medium Non-Essential Amino Acids). Next, slices were mounted using live imaging medium and 200 µm thick iSpacer^®^ (SunJin Lab, Hsinchu, Taiwan). Finally, images were acquired using the 40x magnification objective (NA: 1.1, Immersion: Water) from the Leica TCS SP8 MP system described above. Time lapses of 5-10 min were acquired with a time interval of approximatively 25 sec and a scan format of 512x512 pixels. A brightness/contrast enhancement was performed using imageJ/Fiji.

### Mathematical model of viral-immune interactions in the SGs

For a detailed description of the mathematical model and its parameterization to analyze the infection and immune dynamics in the SG please refer to **Supp. Information – Mathematical model**. In brief, a multi-scale cellular Potts model (CPM) for the SMG was developed using the software Morpheus (http://morpheus.gitlab.io)66. The modeling framework allows us to follow the dynamics and motility of individual cells while accounting for intra- and extracellular processes. Recapitulation of the tissue structure was achieved by segmenting microscopy images of cross sections of the SMG using Ilastik Version1.3.3.pos3 (https://www.ilastik.org)67. Duct structures extracted as binary masks were considered as background and non-duct areas later populated by epithelial cells within Morpheus. The model considers the cellular turnover and infection of immobile epithelial cells, intracellular viral replication within infected cells, as well as export and diffusion of extracellular virus. Endogenous CD4^+^ and M25 CD4^+^ T cells are able to infiltrate the tissue and release IFN_γ_ upon encounter of APC or infected cells dependent on the scenario, thereby inhibiting viral replication within infected cells. All processes of cell proliferation, death, infection and infiltration follow stochastic processes, which were parameterized according to experimental data from the literature or current observations within this study. For processes where reliable measurements are missing, parameter sweeps were performed with model results challenged against the experimental data to evaluate reasonable parameterizations. Sensitivity of results with regard to selected parameter combinations were performed. For computational efficiency, only a small section of 800×800 μm^2^ of the SMG tissue was simulated, with the actually observed 2D-tissue cross sections being 31-34-fold larger (e.g. ∼20-22 mm^2^). Therefore, our simulations generally provide a semi-quantitative representation of the experimentally observed dynamics. Due to the relatively small tissue area simulated, higher infection rates were considered in order to still be able to characterize spatial relationships of infected and immune cells. Otherwise, approximately only 1 in 10 simulations would contain one infected cell around day 8 post infection.

### Statistical analysis

Statistical significance was determined by two-tailed unpaired t test, 2-way Anova with post hoc Tukey’s multiple comparisons test, or two-tailed Pearson correlation using GraphPad Prism (Version 8.2, La Jolla, CA, USA) with *P*<* 0.05, **P*<* 0.01, ***P*<* 0.001 and ****P*<* 0.0001.

## Supporting information

Graphical abstract

## Acknowledgment

We thank Nathalie Oetiker and Franziska Wagen for excellent technical assistance and members of the Oxenius, Joller and Sallusto groups for helpful discussions. We thank the Scientific Center for Optical and Electron Microscopy (ScopeM) at ETH Zurich and the Center for Microscopy and Image analysis (ZMB) at the University of Zurich for their professional support. We very appreciate the important work of our animal facility care takers at ETH Hönggerberg and the very obliging service of Indica Labs providing us with the HALO^®^ software. This work was supported by ETH Zurich and the SNF (IZHRZ0_180552 to AO). FG is supported by the Chica and Heinz Schaller Foundation and member of the IWR. For high-performance computational analyses this work was supported by the state of Baden-Württemberg through bwHPC (MLS-WISO) and the German Research Foundation (DFG) through grant INST 35/1134-1 FUGG.

---

**Supplementary Figure 1.**
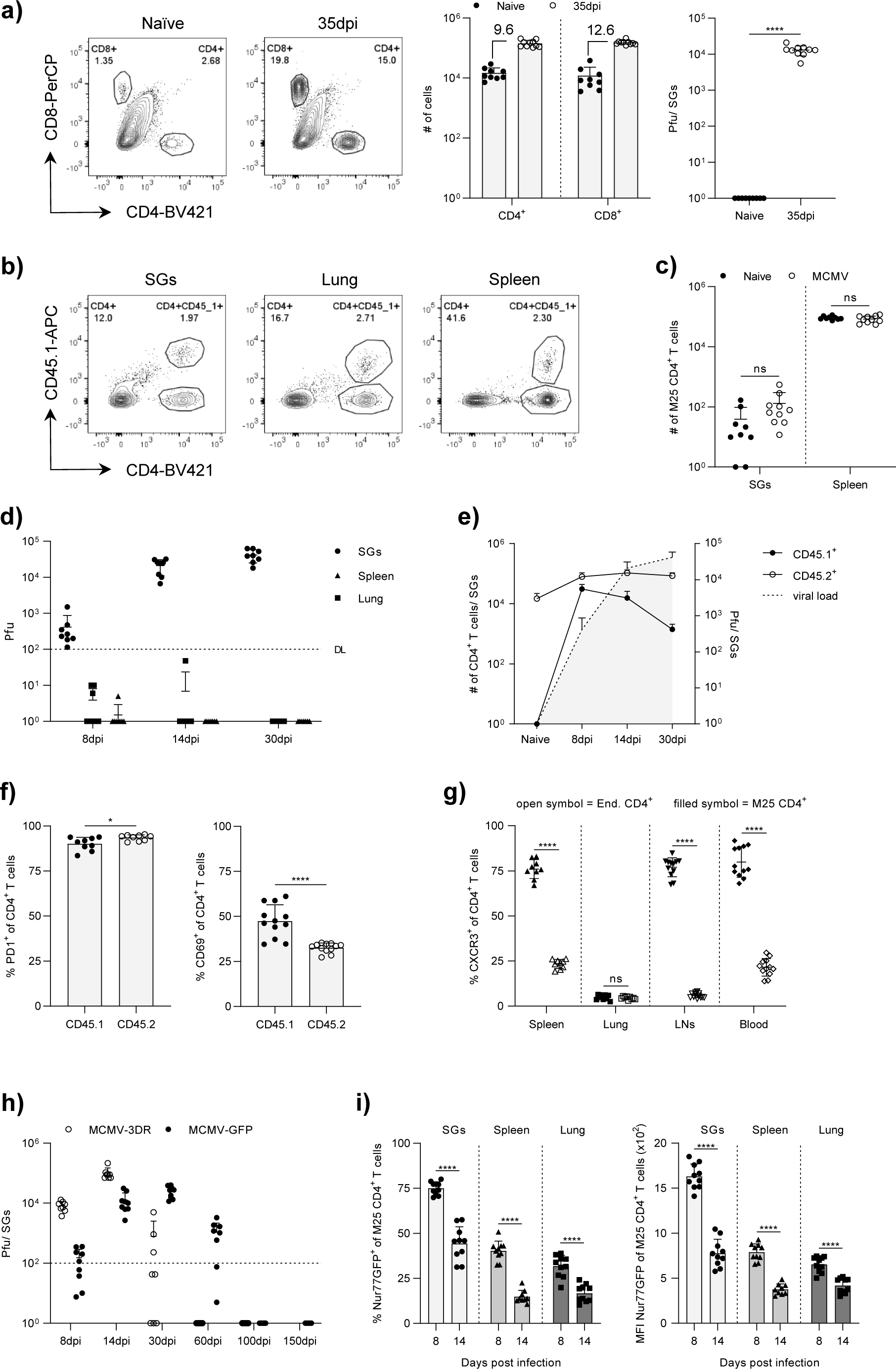
Kinetics and phenotypic characterization of M25 CD4^+^ T cells. **a)** Representative flow cytometry contour plots showing frequency of SG-localized CD4^+^ and CD8^+^ T cells in naïve WT B6 mice and upon MCMV-GFP infection (left) with total T cell counts (middle) and viral loads (right) in the SGs (n = 9 – 10 mice). **b)** Representative flow cytometry contour plots of the SGs, spleens and lungs showing endogenous CD45.2^+^ CD4^+^ T cells and adoptively transferred CD45.1^+^ M25 CD4^+^ T cells. **c)** Total numbers of M25 CD4^+^ T cell counts in the SGs and spleens one week post transfer into naïve or four weeks-infected B6 mice (n = 9 – 10 mice). **d)** Viral loads in the SGs, spleens and lungs at indicated time points post MCMV-GFP infection (n = 8 mice). **e)** Combined kinetics of SGs-infiltrating endogenous CD45.2^+^, adoptively transferred CD45.1^+^ M25 CD4^+^ T cells, and viral burden in the SGs (n = 7 – 9 mice). **f)** Percentage of PD-1^+^ and CD69^+^ CD4^+^ T cells in the SGs (n = 9 mice). **g)** Percentage of CXCR3^+^ CD4^+^ T cells in various organs at day 8 post MCMV-GFP infection (n = 9 – 12 mice). **h)** Kinetics of viral loads in the SGs upon MCMV-3DR or MCMV-GFP infection (n = 8 – 9 mice). **i)** Percentage of Nur77GFP^+^ M25 CD4^+^ T cells (left) and MFI of Nur77GFP on M25 CD4^+^ T cells (right) in SGs, spleens and lungs at day 8 and 14 post MCMV-3DR infection (n = 9 – 10 mice). Data in **a** and **c – i** are shown as mean + SD pooled from two independent experiments. Each symbol represents an individual mouse. Statistical significance was determined using single (**a and f**) or multiple (**c, g and i**) unpaired two-tailed t test. ****P<0.0001, ns = not significant.

**Supplementary Figure 2.**
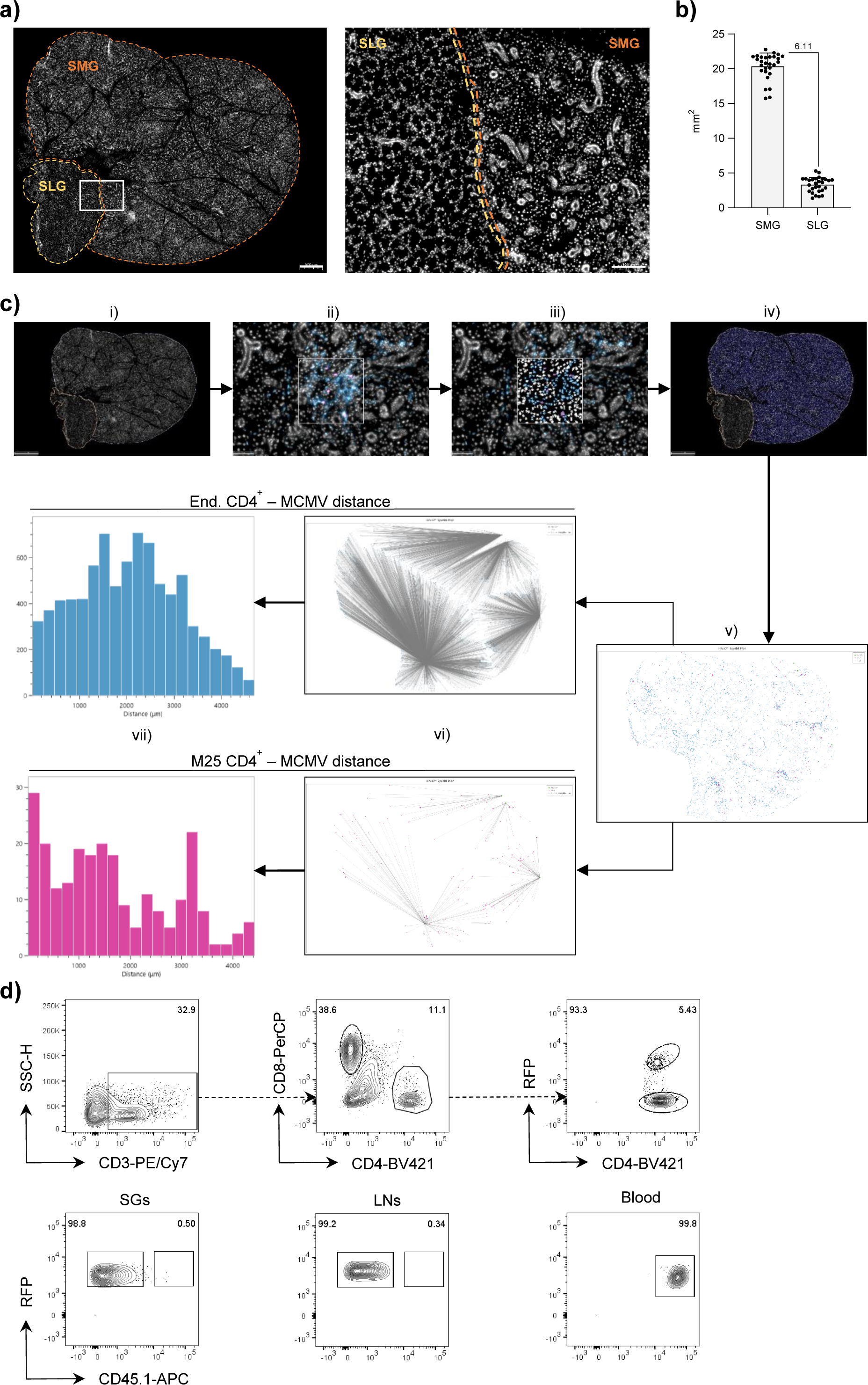
Spatial localization of endogenous CD4^+^ and M25 CD4^+^ T cells in the SGs. **a)** Example of the right SG lobe with magnified region; perimeters of the submandibular and sublingual gland, SMG and SLG respectively, are indicated by dotted lines. **b)** Area of SMGs and SLGs. **c)** Image processing procedure: i) Tissue Annotation (i.e. SMGs and SLGs), ii) unmodified, original FOV within the SMGs and iii) after applying cell segmentation and phenotypic characterization (HighPlex FL module), iv) application of the HighPlex FL module to the whole annotation layer (i.e. SMGs), v) “Reduced” image consisting of extracted endogenous CD4^+^ and M25 CD4^+^ T cells from total cells together with infection foci, vi) Nearest neighbor analyses between endogenous CD4^+^ or M25 CD4^+^ T cells and infection foci and vii) representative histogram of overall distances between endogenous CD4^+^ and M25 CD4^+^ T cells and infection foci. **d)** I.v. labeling of M25 CD4^+^ T cells in various organs. MACS purified, RFP expressing M25 CD4^+^ T cells were adoptively transferred into naïve WT B6 one day prior MCMV-GFP infection. At day 8 post MCMV-GFP infection, 3 μg of anti-CD45.1 antibody was injected i.v. 3 min prior harvest. M25 CD4^+^ T cells were identified by the RFP signal. Upper row: Gating strategy, lower row: Representative flow cytometry contour plots of SGs, SG draining LNs and blood. Plots are representative of n = 5 mice from one experiment. Scale bar in **b** = 500 μm (left) and 100 μm (right).

**Supplementary Figure 3.**
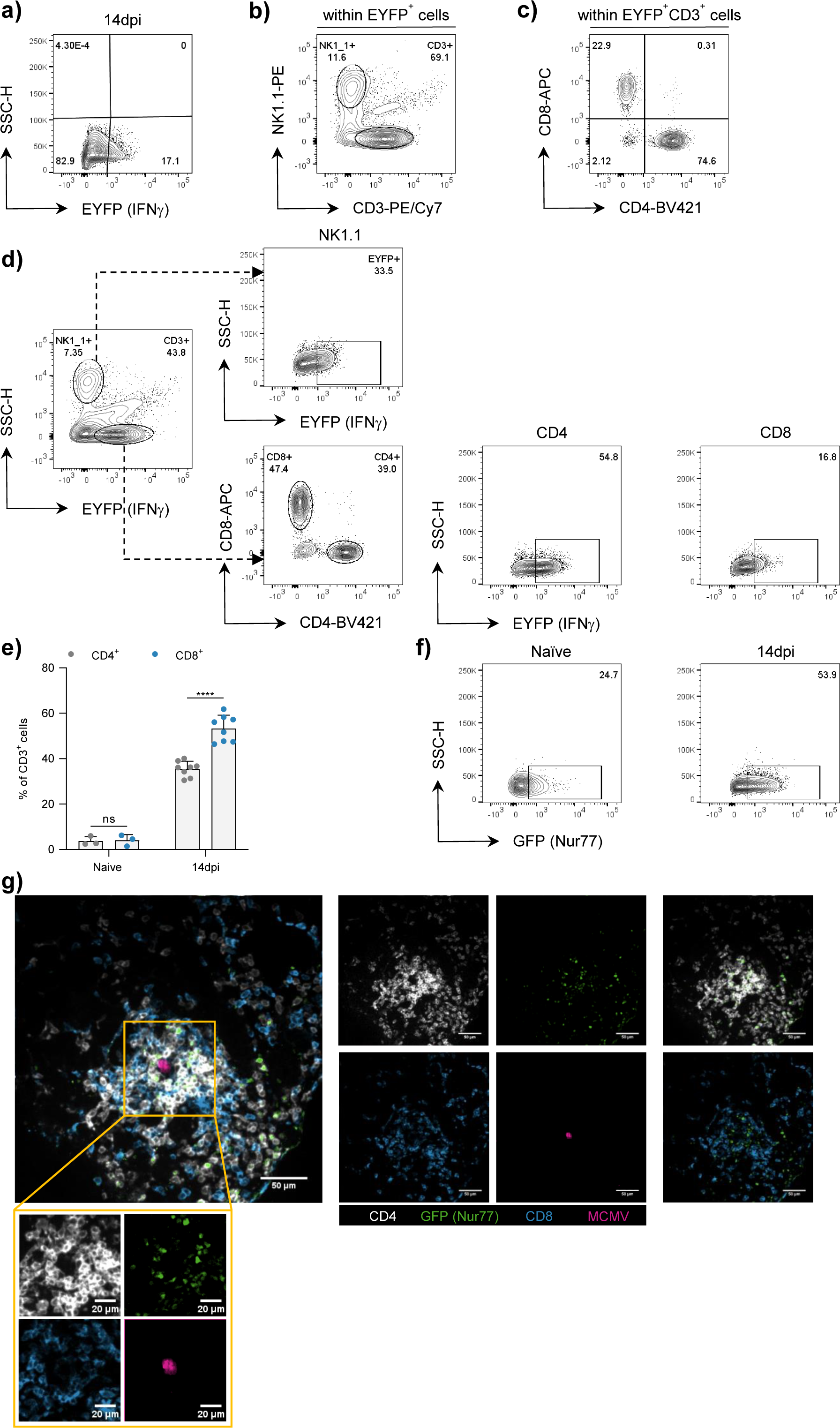
IFN_γ_ production by CD4^+^ T cells in MCMV-3DR-infected SGs. **a–c)** Representative flow cytometry contour plots showing gating strategy. **a)** Frequencies of EYFP^+^ cells upon MCMV-3DR infection. **b and c)** Further subgating of the EYFP^+^ cell fraction. **d)** Representative flow cytometry contour plots of MCMV-3DR infected SGs showing gating strategy to determine the percentage of EYFP^+^ cells within CD4^+^, CD8^+^ and NK1.1^+^ cells. **e)** Percentage of CD4^+^ and CD8^+^ T cells within CD3^+^ cells in naïve and MCMV-3DR infected Great mice. **f)** Representative flow cytometry contour plots of Nur77GFP expression in SG-residing CD4^+^ T cells in naïve and MCMV-3DR infected Nur77GFP mice. **g)** Example of a four color FOV of a Nur77GFP mouse-derived SG 14 days post MCMV-3DR infection (left). Single color channel images (middle), merged two color CD4/GFP (right, upper row) and CD8/GFP (right, lower row) images. Magnified region (yellow frame) region split in four single color channel images. Data in **e)** are shown as mean + SD of n = 3 and n = 8 infected Great mice pooled from 2 independent experiments. Statistical significance in **e)** was determined using multiple unpaired two-tailed t test. ****P<0.0001, ns = not significant. Scale bar in **g** = 50 μm or 20 μm (magnified region).

**Supplementary Figure 4.**
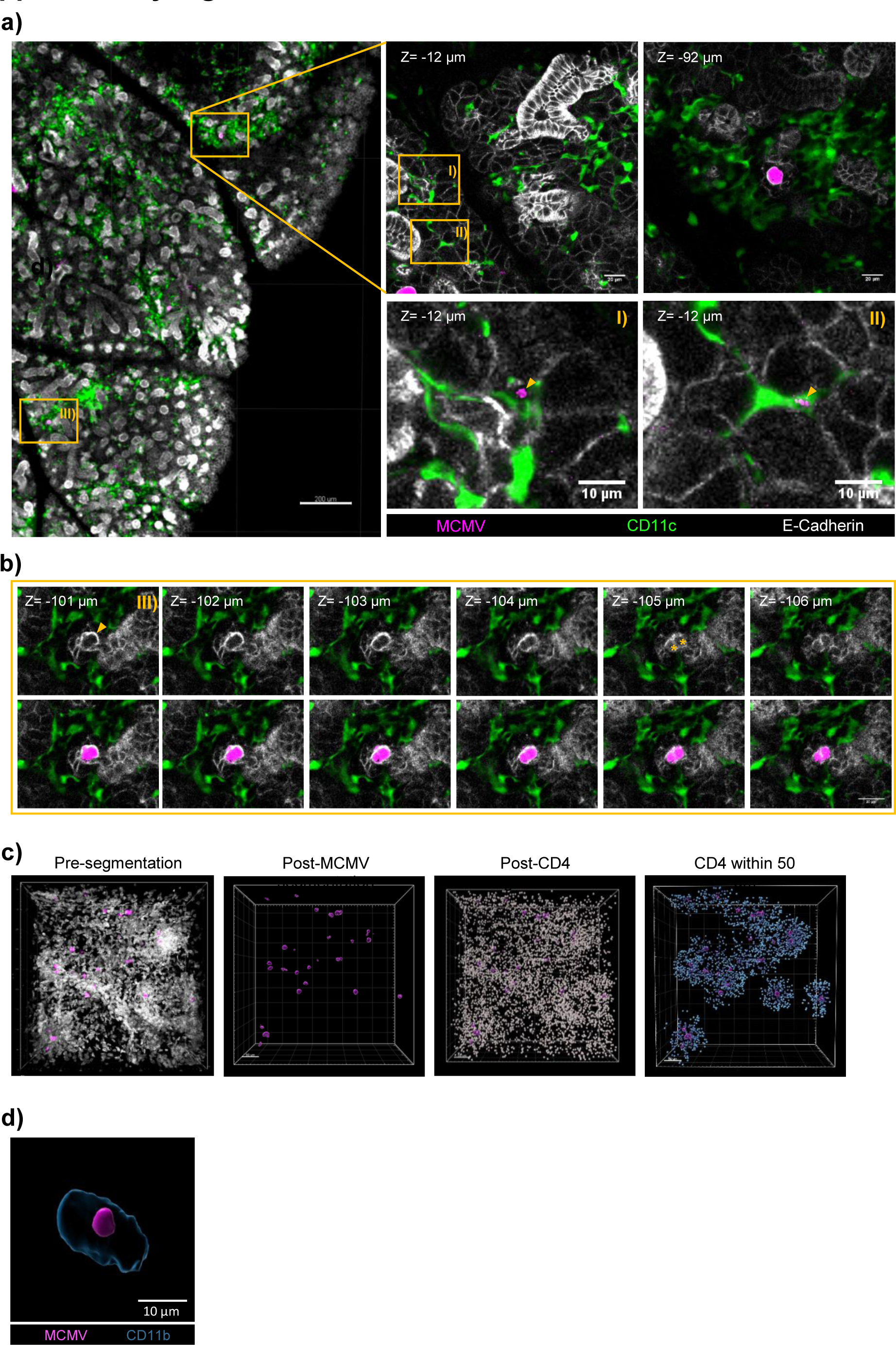
Processing and assignment of MCMV-3DR signals in the SGs. **a)** Example of a three color FOV of a CD11cYFP-Prox1morange mouse-derived SG 14 days post MCMV-3DR infection (left, scale bar = 200 µm) with magnified regions (right, scale bar = 20 µm (top) and 10 µm (bottom)) at indicated z-positions. **b)** Magnified region of **a** (yellow frame: III) at sequential z-planes without (upper row) and with (lower row) MCMV-3DR encoding mcherry signal (scale bar = 20 µm). Yellow arrowheads in **a** and **b** indicate engulfment of possible remnants of previously infected cells in CD11c^+^ cells and large actively infected E-Cadherin^+^ cells, respectively. Yellow asterisks in **b** show two neighboring MCMV-3DR infected E-Cadherin^+^ cells. **c)** Segmentation process for spatial analysis. From left to right: Unprocessed 3D reconstruction, post-segmentation of MCMV signal as surfaces, post-segmentation of the CD4 signal as spots and CD4^+^ T cells within 50 µm to next site of viral replication. Scale bar = 50 µm. **d)** 3D reconstructions of MCMV-3DR-associated mcherry signal within a CD11b^+^ cell 8 dpi. Scale bar = 10 µm.

**Supplementary Figure 5.**
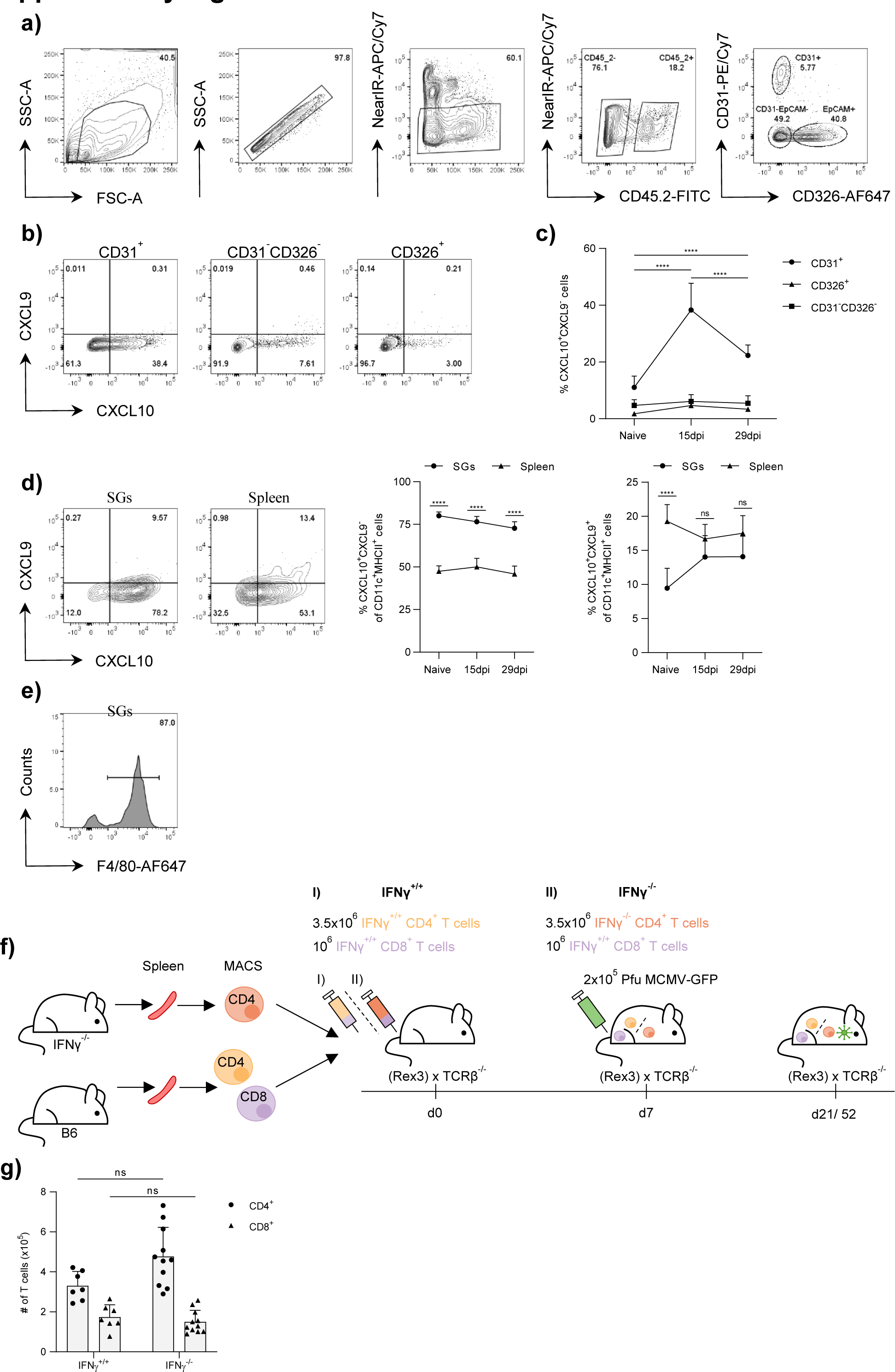
CXCL9 and CXCL10 expression and their role in MCMV-infected SGs. **a and b)** Representative flow cytometry contour plots showing gating strategy of non-hematopoietic cells **(a)** and CXCL9 and CXCL10 expression levels **(b)** in defined non-hematopoietic cell compartments 15 days post MCMV-GFP infection **c)** Percentage of indicated CXCL10^+^ non-hematopoietic cells at indicated time points. **d)** Representative flow cytometry contour plots (left) and percentages (middle & right) of CXCL9 and CXCL10^+^ CD11c^+^MHCII^+^ cells at indicated time points post MCMV-GFP infection in the SGs and spleen. **e)** Representative histogram of F4/80 expression levels in CD11c^+^MHCII^+^ cells of the SGs 15 days post MCMV-GFP infection. **f)** Experimental approach. T cells from spleens of naïve WT B6 (IFN_γ_^+/+^) and IFN_γ_KO (IFN_γ_^-/-^) mice were purified using anti-CD4 or anti-CD8α MACS beads. 3.5x10^6^ IFN_γ_ competent (IFN_γ_^+/+^) or IFN_γ_ deficient (IFN_γ_^-/-^) CD4^+^ T cells and 10^6^ IFN_γ_ competent CD8^+^ T cells were adoptively co-transferred into naïve Rex3 x TCRλ3^-/-^ or TCRλ3^-/-^ mice seven days prior MCMV-GFP infection. Flow cytometric and microscopic analyses of CXCL9 and CXCL10 expression were performed 14 dpi, viral burden was evaluated 14 and 45 dpi. **g)** Total numbers of CD4^+^ and CD8^+^ T cells in the SGs 14 dpi. Data in **c** and **d** are shown as mean + SD of n = 6 - 8 Rex3 mice pooled from two independent experiments. Each symbol in **c** and **d** represents the mean of pooled mice. Statistical significance was determined using 2-way Anova with post hoc Tukey’s multiple comparisons test (**c**) or multiple unpaired two-tailed t test (**d & g**). ****P<0.0001, ns = not significant.

**Supplementary Figure 6.**
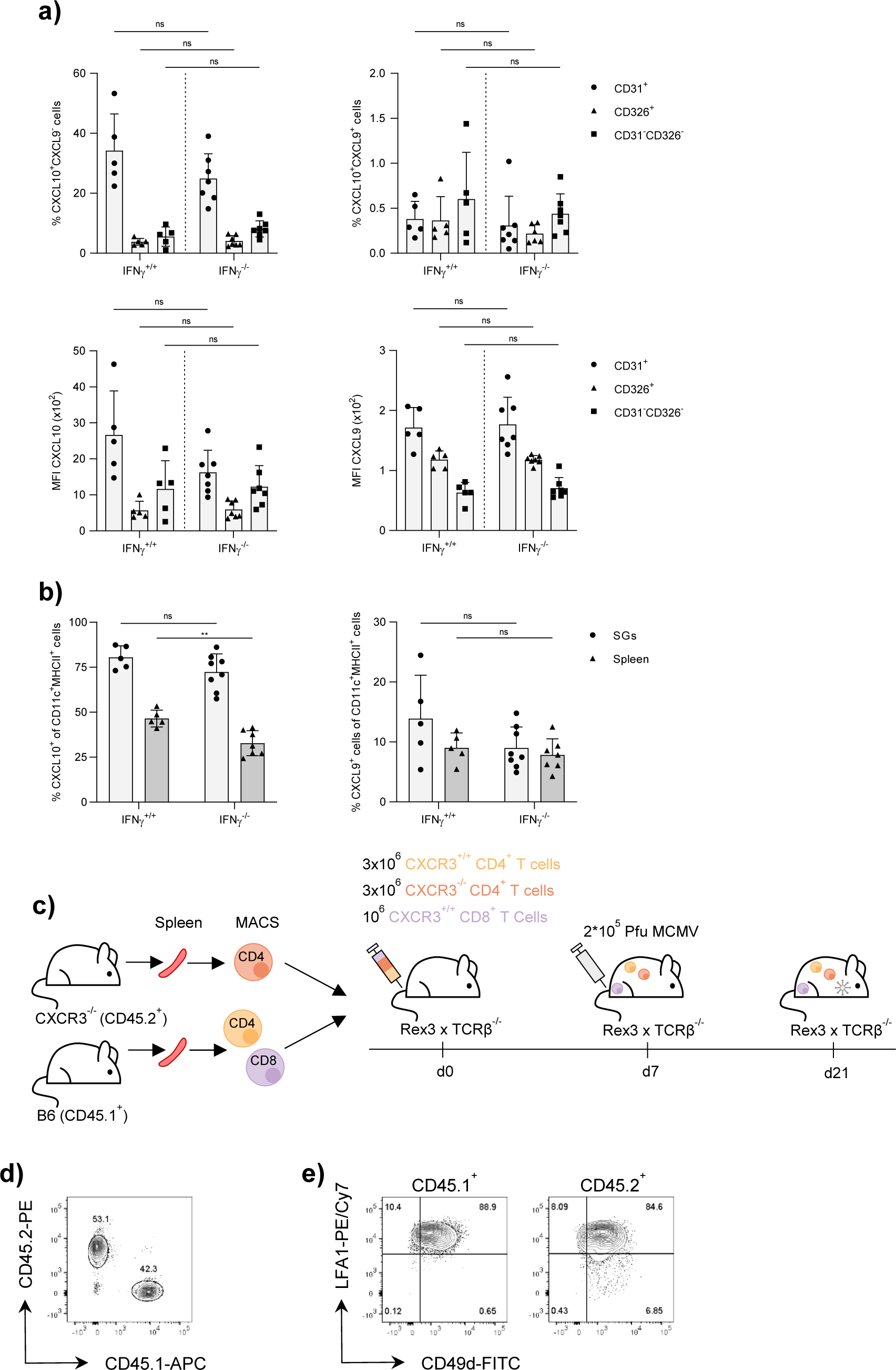
CXCR3-mediated recruitment of T cells to sites of CXCL9 and CXCL10 expression in MCMV-infected SGs. **a)** Percentage (upper row) and MFI (lower row) of CXCL9 and CXCL10 expression in indicated non-hematopoietic cell types in the SGs 14 dpi. **b)** Percentage of CXCL9^+^ and CXCL10^+^ CD11c^+^MHCII^+^ cells in the SGs and spleen 14 dpi. **c)** Experimental approach. T cells from spleens of naïve WT B6 (CD45.1^+^, CXCR3^+/+^) and CXCR3^-/-^ mice (CD45.2^+^) were purified using anti-CD4 or anti-CD8α MACS beads. 3.*10^6^ CXCR3 competent (CD45.1^+^CXCR3^+/+^) and CXCR3 deficient (CD45.2^+^CXCR3^-/-^) CD4^+^ T cells, together with CXCR3 competent CD8^+^ T cells, were adoptively co-transferred into naïve Rex3 x TCRλ3^-/-^ mice seven days prior MCMV infection. Flow cytometric and microscopic analyses were performed 14 dpi. **d)** Representative flow cytometry contour plots of CD45.1^+^(CXCR3^+/+^) and CD45.2^+^ (CXCR3^-/-^) CD4^+^ T cells and **e)** CD49d^+^LFA-1^+^ CD4^+^ T cells in the SGs 14 dpi. Data in **a & b** are shown as mean + SD of n = 5 - 8 Rex3 x TCRλ3^-/-^ mice pooled three independent experiments. Graphs in **d)** and **e)** are representative of n = 4 mice from one independent experiment. Each symbol in **a** and **b** represents an individual mouse. Statistical significance was determined using multiple unpaired two-tailed t test (**b**), or 2-way Anova with post hoc Tukey’s multiple comparisons test (**c & d**). *P<0.05, ns = not significant.

**Supplementary Figure 7.**
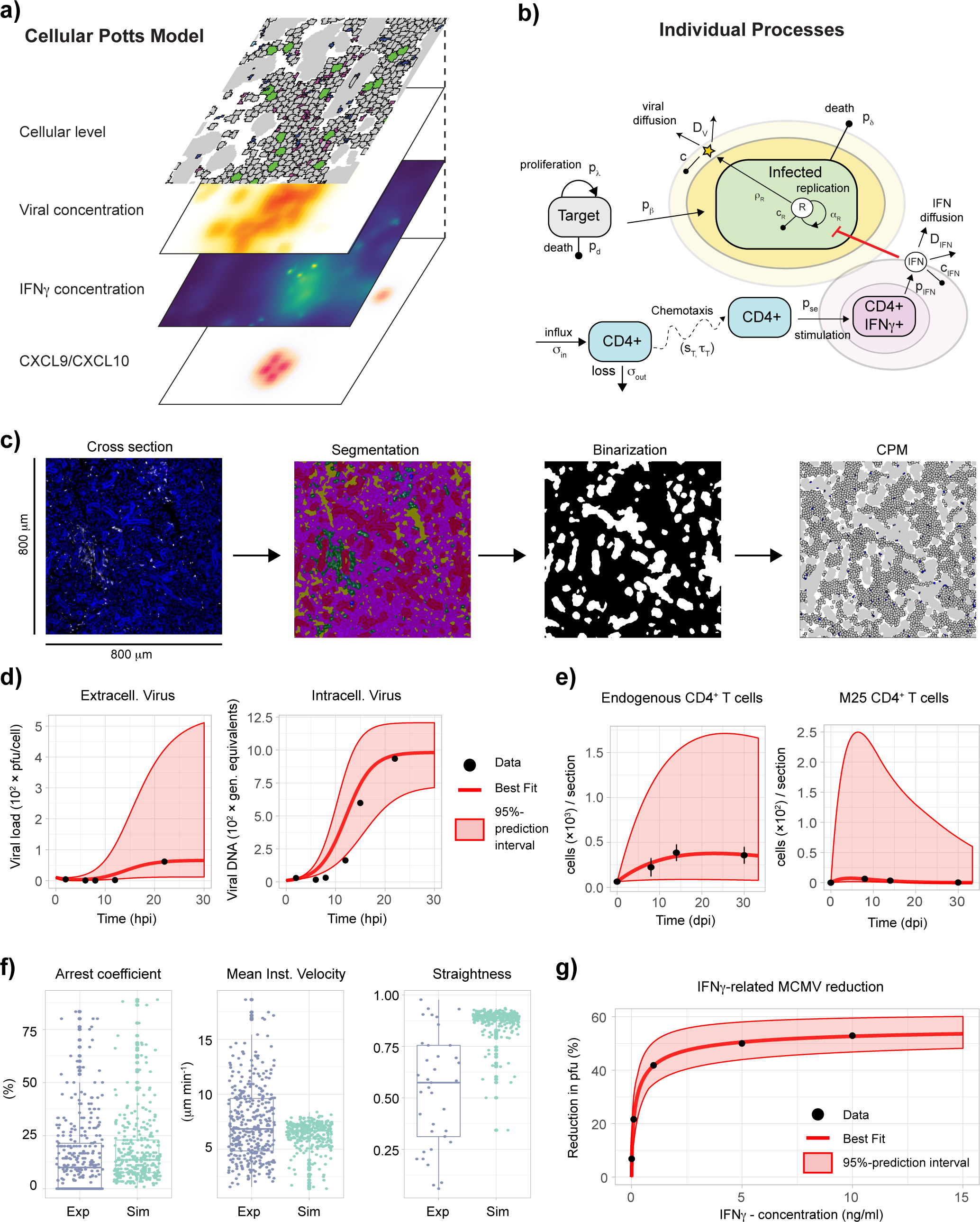
Mathematical modeling of viral-immune interactions in the SGs. **a)** Sketch of the structure of the multi-scale cellular Potts model used to simulate viral-immune interaction in the SMG. Different interconnected layers account for the spatiotemporal dynamics of individual cells in interaction with CXCL9/CXCL10, IFN_γ_, and extracellular MCMV concentrations. **b)** Individual processes considered within the CPM that describe the turnover and infection of epithelial cells, intracellular viral replication, as well as the motility and stimulation of CD4^+^ T cells, and diffusion of virus and IFN_γ_. Individual parameters are explained in detail within the **Supplemental Information - Text S1**. **c)** Representation of the ductal structure within the simulations by using 800×800 μm^2^ areas selected from SMG cross section images that were segmented to distinguish ductal cells (*red*), M25 (*blue*) and endogenous (*green*) CD4^+^ T cells, cell free areas (*yellow*), and remaining cells (purple) using *HALO* and *ilastik* software. Segmented images were converted to binary images to distinguish between ductal (*white*) and non-ductal (*black*) areas, with non-ductal areas later populated by epithelial cells within the CPM. **d)** Experimental data digitalized from (Misra et al., 1977) and best fit to describe MCMV viral replication dynamics based on Eq. (8). **e)** Infiltration dynamics of endogenous (upper panel) and M25 CD4^+^ T cells (lower panel) using Eq. (9) and experimental data (see **Text S1**). **f)** CD4^+^ T cell motility dynamics as predicted by fitting the CPM to the data of (Stolp et al., 2020) on CD8^+^ T cell dynamics in the SG. The distributions of the arrest coefficient, mean instant velocity and straightness for the data (*blue*) and model predictions (*cyan*) following 300 T cells over 1h are shown, with model predictions based on the best parameterization obtained by the FitMultiCell approach. **g)** Dose-response curve for the effect of IFN_γ_ on MCMV replication efficiency parameterized based on Eq. (10) and the data of (Lucin et al., 1994). Panels **(d)**, **(e)**, and **(g)** indicate experimental data (*black dots*), best model fits (*red line*) and 95%-prediction bands (*red shaded area*). All individual parameter estimates are shown in **Text S1**.

**Supplementary Video 1. 3D visualization of CD4^+^ T cells and MCMV-3DR-associated mcherry signals (sites of viral replication and small remnants).** 3D reconstruction of CD4^+^ T cells (white) and MCMV-3DR-associated mcherry signals (magenta) in the SMG at 8 dpi. Small mcherry volumes represent remnants or apoptotic bodies and large mcherry volumes sites of infectious virus production.

**Supplementary Video 2. Quantitative analysis of CD4^+^ T cells within 50 µm to each MCMV- 3DR-associated mcherry signal.** Successive 3D processing of mcherry-expressing MCMV-3DR (magenta) as surfaces and CD4^+^ T cells (white) as spots with further thresholding of these spots within 50 µm (cyan) to each MCMV-3DR-associated mcherry signal in the SMG. Major tick intervals = 50 µm.

**Supplementary Video 3. *Ex vivo* live imaging of CD11c^+^ cell phagocytosing MCMV-3DR- derived small remnants.** Times series of CD11c^+^ cell (green) phagocytosing surrounding MCMV- 3DR-associated remnants (magenta) at 9 dpi. Scale bar = 3 µm. Time in hours: minutes: seconds: milliseconds.

**Supplementary Video 4. 3D reconstruction of accumulating CD11c^+^ cells and CD4^+^ T cells.** 3D processing of MCMV-3DR-associated mcherry signals (magenta), CD4^+^ T cells (white) and CD11c^+^ cells (green) as surfaces in the SMG at 14 dpi. Zoom-in on small remnants within CD11c^+^ cells.

## Supplemental Information Text S1

### Mathematical model of MCMV infection and immune dynamics in the salivary glands

We developed a multi-scale mathematical model to study the spatiotemporal dynamics of MCMV infection and CD4^+^ T cell dynamics in the salivary glands (SG). The model follows the dynamics and fate of individual cells while accounting for intra- and extracellular processes, such as intracellular viral replication and extracellular viral diffusion, respectively. It is built as a cellular Potts Model (CPM) using the software Morpheus (http://morpheus.gitlab.io;(Starruss et al., 2014)), which provides a 2D-representation of the observed tissue structure and accounts for individual cell morphology, motility and their spatial location. A CPM is a lattice-based model with individual cells defined by several connected lattice sites (Glazier and Graner, 1993; Graner and Glazier, 1992). Each cell is characterized by a specific cell type, as well as a unique identifier w and the position within the grid, (*i,j*). The motility and dynamics of individual cells is then determined by stochastic events that regulate the interaction between grid sites (Glazier and Graner, 1993). Our model considers the turnover and infection of epithelial cells, the diffusion and clearance of extracellular virus, as well as the infiltration, migration and locally confined release of IFN_γ_ of endogenous and M25-specific CD4^+^ T cells. A sketch of the model and its structure is shown in **Figure S7a,b**. A detailed description of the individual model components, as well as their parameterization based on own experimental data and data from the literature is given in the following.

### The multi-scale mathematical model

#### Epithelial cells and viral replication

The multi-scale model provides a 2D-representation of the tissue structure of the submandibular salivary glands (SMG) as observed in the experimental cross sections. Epithelial cells, which form the structure of the solid tissue, are immobile, and die and proliferate with probability *p*_λ_ and *p*_d_, respectively. Hereby, *p*_A_ is defined by

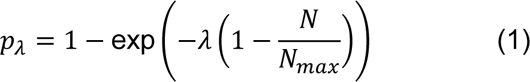

with *11* defining the maximal proliferation rate, *Nmax*, the maximal cell density and *N* the current number of epithelial cells to ensure tissue homeostasis in response to cell turnover or tissue damage. Uninfected epithelial cells, i.e., target cells, get infected by MCMV dependent on the viral concentration, *V*, at the location of the cell. The probability of infection, *p*_/3_, is given by

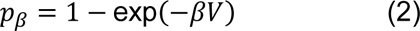

Hereby, the parameter *λ3* defines a scaling factor, which determines transmission of infection dependent on the extracellular viral concentration. Upon infection, epithelial cells will double in size (Munger et al., 2006) and start replication of intracellular viral RNA, *R*, which is modeled by a system of ordinary differential equations following logistic-growth with

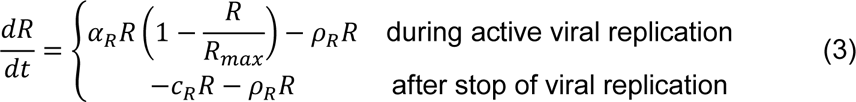

Hereby, *Rmax* defines the maximal carrying capacity of the cell, and *αR*, *πR* and *c*_R_ the net-replication, viral export and intracellular viral clearance rate, respectively. The stop of viral replication, as e.g. mediated by IFN_γ_, is represented by the switch to the second case in Eq. (3) in which the virus gets degraded with rate *c*_R_ and is continued to be exported until intracellular virus vanishes (see also (Rand et al., 2012)). As MCMV suppresses cell cycle progression (Marcinowski et al., 2012), cell division of infected cells is not considered within the simulations,while death occurs with probability *p*_8_.

Once exported, extracellular virus diffuses through the tissue with diffusion coefficient *DV*. Thus, the concentration of extracellular virus is modeled by a reaction-diffusion system given by

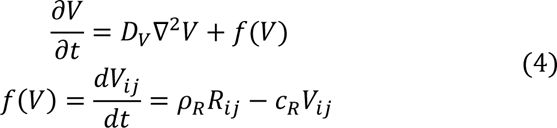

Hereby, *f(V)*, describes viral export, *πR*, and viral degradation, *c*_R_, at each specific lattice site *(i,j)* within the grid, with *Rij* = 0 for all lattice sites that belong to uninfected cells. To determine the probability of an uninfected cell, χο, to get infected (Eq. (2)), the viral concentration is averaged over all lattice sites that belong to the uninfected cell, i.e, 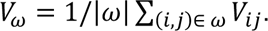

Local concentrations of IFN_γ_ can inhibit viral replication in infected cells with probability *p*_IFNy_ defined by

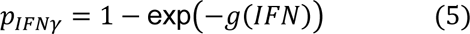

with *g*(*IFN*) determining the dose-response relationship for IFN_γ_ given by

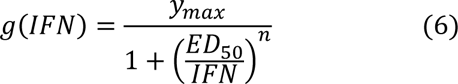

Hereby define *y*_max_ the maximal inhibition efficacy, *ED*_SO_ the effective dose at which 50% of cells are inhibited, and *n* a Hill-coefficient. Upon inhibition of intracellular viral replication, the concentration of intracellular viral RNA will follow the second case of Eq. (3). As the effect of IFN_γ_ on the replication rate of MCMV infected cells was shown to be reversible (Kropp et al., 2011; Presti et al., 1998), infected cells can loose the replication inhibition without maintained IFN_γ_ signaling. An infected cell is considered as an active, MCMV replicating cell (MCMV^+^) and, thus, visible by light microscopy, if the intracellular viral load R is larger than a threshold of 100 (R>100).

#### CD4^+^ T cells and IFN**γ** release

In contrast to epithelial cells, endogenous and M25 CD4^+^ T cells are considered as motile cells that enter and leave the tissue with time-dependent influx and efflux rates, σ_in_ and σ_out_. Within the tissue, CD4^+^ T cells follow persistent motion modeled with persistent strength sT and decay time 1T. The movement of CD4^+^ T cells can be influenced by chemokine fields, e.g. CXCL9 and CXCL10 chemotactic gradients as mediated by antigen presenting cells (APC). The attraction of cells is defined by the chemotactic strength, *CTX*, that determines the strength at which the gradient can deviate the direction of the CD4^+^ T cell. Stimulation of CD4^+^ T cells leading to release of IFN_γ_ depends on interaction of CD4^+^ T cells with APC or infected cells dependent on the scenario (see below). When stimulated, the CD4^+^ T cell becomes temporally immobile and will secrete IFN_γ_ with rate σ_IFN_. Analogously to the extracellular virus, the dynamics of the IFN_γ_ concentration within the system is then described by

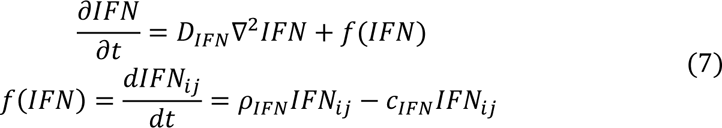

Hereby, D_IFN_ defines the 2D-diffusion coefficient of IFN_γ_, and *f*(*IFN*) describes the release, *ρ*_IFN_, and extracellular clearance, *c*_IFN_, of IFN_γ_ at each specific lattice site *(i,j)*. To calculate the probability of inhibiting viral replication in infected cell *χοI* according to Eq. (3) and (4), the IFN_γ_ concentration is averaged over all lattice sites that belong to an infected cell, i.e., 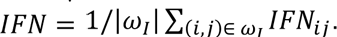

### Parameterization and Model adaptation

The individual components and processes of the model were parameterized based on own experimental data and data from the literature. For some of the processes where reliable measurements are missing, parameter sweeps were performed with model results challenged against the experimental data to evaluate reasonable parameterizations. Sensitivity of results with regard to selected parameter combinations were performed.

#### SGM tissue structure

Recapitulation of the tissue structure was achieved by segmenting microscopy images of cross sections of the SMG using Ilastik Version1.3.3.pos3 (https://www.ilastik.org, (Berg et al., 2019)). Duct structures were extracted as binary masks and loaded as background into Morpheus. The images were then populated by immobile epithelial cells representing acinar glandular epithelial cells as the main target cells for MCMV (Fox, 2006; Walton et al., 2011). Duct areas were left empty (**Figure S7c**). The spatial localization of immune and infected cells was measured on 20-22 m^2^ cross sections, which is an unfeasible system size for the simulations. For computational efficiency, only small sections of 800×800 μm^2^ of the SMG tissue were considered, with the actually observed 2D-tissue cross sections being 31-34-fold larger (e.g. ∼20-22 mm^2^). Simulations were run with a scaling factor of 1 pixel (px) = 2 μm to allow simulation of larger areas. The cell area of target and infected cell was determined using microscopy images of the SG cross sections with labelled MCMV-GFP infected cells. The average diameter of infected cells was determined as ∼17.54 μm leading to an average area of 241.6 μm^2^. Target cells, i.e., uninfected epithelial cells, were set to half of the area of infected cells (Kwak et al., 2016; Munger et al., 2006), i.e., 120.8 μm^2^ **(Table A)**.

At the beginning of a simulation, ∼2500 target cells were initialized within the grid defining the maximal population capacity of the system *Nmax*. This corresponds to the average area of ∼46% that was covered by non-immune and non-duct cells within the observed cross sections. With epithelial cells usually having a relatively long lifespan of 2-4 months (Aure et al., 2015; Schwartz- Arad et al., 1988), the probability of uninfected epithelial cells to die, *p*_d_, is ρ-distributed with scale and shape parameters determined by kT=40 and 8T=55 leading to an expected lifespan of ∼3 months. With MCMV-infected cells observed to remain viable for 36-48h (Fox, 2006) cell death of infected cells, *p*_8_, was modeled analogously according to ρ(kI =0.25,8I =170).

#### Intracellular viral replication

The dynamics of intracellular viral replication and export as described in Eq. (3) was parameterized using *in vitro* experimental data on the viral synthesis in MCMV infected mouse embryo fibroblasts (MEF)(Misra et al., 1977). In this experiment, MEFs in cell culture were infected with MCMV and the number of genome equivalents and pfu per cell were determined at frequent time points in between 0 and 22 hours post infection **(Figure S7d)**. The data allowed us to simultaneously estimate the net-viral replication rate, *α*_R_, extracellular viral degradation, *c*, and viral export, *ρ*_R_, using a single coherent data set. To analyze the data and estimate these parameters, the system in Eq. (3) was extended by

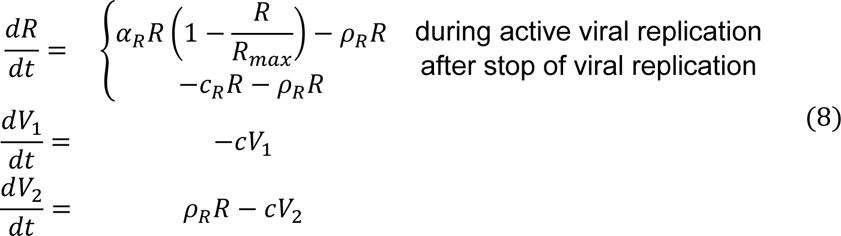

which distinguishes between the extracellular viral load applied to the system at the start of the experiment, *V*_1_, and the newly produced virions by the infected cells, *V*_2_. Eq. (8) was fitted to the data using *V* = *V*_1_ + *V*_2_. Based on these analyses we estimated a net viral replication rate of *α*_R_=0.41 h^-1^ (0.38, 0.49), a maximal carrying capacity of *R*_max_= 1060 genome equivalents (1020,1260), a viral export rate of *ρ*_R_= 0.03 pfu/gen. equivalent h^-1^ (0.02, 0.11) and a viral degradation rate of *c* = 0.45 h^-1^ (0.20, 1.65), with numbers in brackets indicating 95%-confidence intervals of estimates. As the rates of intracellular viral replication and degradation could not be determined separately based on these experimental data, the rate of intracellular RNA degradation, *c*_R_, was set to the extracellular degradation rate, *c*_R_ = *c*. The experimental data with model predictions and all parameter estimates are shown in **Figure S7d** and **Table A**, respectively. For technical details on the fitting procedure please see section ***Parameter fitting and estimation procedures***. While the model predicts saturation of the viral concentrations ∼24 h corresponding to the assumed length of the replication cycle of MCMV (Misra et al., 1977), it slightly underestimates the lag time before the start of rapid viral replication leading to earlier viral export.

#### MCMV spread and cell proliferation

There are three factors that influence the degree and extend of MCMV spread and tissue pathology within the salivary glands in the absence of any immune response: This includes (i) viral spread, as defined by the scaling factor λ3 regulating the infection probability, *p*_/3_, dependent on the extracellular viral concentration, *V* (Eq. (2)), (ii) extracellular viral diffusion defined by the diffusion coefficient *D*_v_ (Eq. (4)), and (iii) tissue regeneration determined by the cell proliferation parameter 11 (Eq. (1)). Individual assessment of the parameters λ3 and 11 is generally difficult due to their strong interdependency and the lack of appropriate experimental data. Therefore, we performed combined sweeps of λ3 and 11 to identify appropriate parameter combinations. Model simulations without the presence of immune responses were challenged against the following observations and assumptions to restrict parameter values:

1. As our data indicate high infection levels at 8 dpi, parameter λ3 should be chosen to allow a considerable spread of the infection within the tissue already at 8 dpi. This is used to set a lower bound for λ3.
2. Even without the presence of CD4^+^ T cell immunity, we assumed that the virus is incapable of leading to the complete lysis of large tissue sections at 8 dpi. These comparisons allowed us to determine an upper boundary to λ3 and a lower bound to 11.
3. MCMV indicates local spread within tissue (Wirtz et al., 2008) leading to infectious foci. This observation was used to balance the relationship between λ3 and *D*_v_ that regulates the extent of viral spread.

Previous measurements indicate a viral diffusion coefficient for MCMV in between *D*_v_ = 4.4 μm^2^/s - 6.4 μm^2^/s (Dick et al., 2015). However, given the long time period (∼60 days) required for our simulations, we used an effective viral diffusion coefficient of 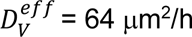 in our model, which resulted in locally confined virus fields around the infected cell. As the choice of 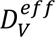 is compensated by the choice of the unknown scaling parameter λ3, this does not affect our analyses, but substantially increases computational efficiency. Otherwise simulation time would increase by ∼100-fold, making it unfeasible for our analyses. Accounting for the observations in (1.) - (3.) mentioned above, we finally set 11= 0.204 h^-1^ and λ3= 1.5×10^-3^ pfu^-1^h^-1^ to scale tissue regeneration and infection spread in our simulations. With these parameters we obtained preferential local spread of the infection as observed in (Wirtz et al., 2008), but also seeding of new infections at distant locations due to viral diffusion. In general, the maintenance of the tissue, as well as the dynamics of infection was sensitive to the choices of 11 and λ3. More detailed measurements will be needed to appropriately curtail both values for future analyses.

#### CD4^+^ T cell infiltration dynamics

In agreement with the experimental data, our model distinguishes between endogenous and M25 CD4^+^ T cells. Both cell types are modeled with a targeted T cell area of ∼50 μm^2^ corresponding to a T cell diameter of ∼8 μm which is at the lower bound of average T cell sizes. To parameterize the infiltration dynamics of CD4^+^ T cells into the SMG during the time course of the infection, we used our experimental measurements. The numbers of endogenous and M25 CD4^+^ T cells were determined at 0, 8, 14 and 30 dpi for the total SMG, and for specified cross sections at 8 dpi. Based on the ratio of CD4^+^ T cells on the cross sections and the total SMG at 8 dpi, we calculated the expected density of endogenous and M25 CD4^+^ T cells in a defined area of 800×800 μm^2^ as used for our simulations **(Table B)**. The population dynamics of CD4^+^ T cells in the SMG can then be described by the following system of ordinary differential equations to determine the net-flux and loss of cells in and out of the SMG,

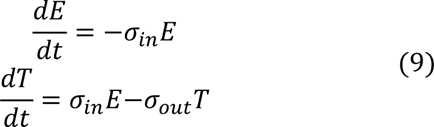

Hereby, *E* and *T* define the concentration of CD4^+^ T cells outside and within the SMG, respectively, and σ_in_ and σ_out_ the rate of influx and loss of cells. Eq. (9) was fitted separately for endogenous and M25 CD4^+^ T cells based on the experimental data shown in **Table B**. Best fits and 95%- confidence intervals are shown in **Table A** with experimental data and model predictions shown in **Figure S7e**. All parameters for endogenous cells were identifiable, while low cell counts slightly impaired parameter identifiability for M25 CD4^+^ T cells.

The estimates were then used to parameterize the appearance and loss of endogenous and M25 CD4^+^ T cells in our CPM. At each time step, a total number of σ_in_*E*(*t*) cells was randomly placed on the grid to simulate CD4^+^ T cell infiltration. In addition, a CD4^+^ T cell was removed from the tissue with probability *p*_out_ = 1 − exp(-σ_out_). For technical reasons, CD4^+^ T cells were initialized at the positions of ductal structures and moved into the tissue from these points of entry. To guide this process, a gradient from the center to the borders of the duct structures was introduced. CD4^+^ T cells moved against the gradient and into the tissue with a chemotactic strength set to *CTX*_duct_ = 1, which resulted in a homogeneous distribution of the CD4^+^ T cells in the grid. Larger values would result in strong borders between the acinar and ductal areas and inhomogeneous distribution of the CD4^+^ T cells over the grid.

#### CD4^+^ T cell motility within the SMG

Parameters to describe CD4^+^ T cell motility within the SMG were determined using data from two-photon intravital microscopy analyses on the dynamics of CD8+ T cells in the SMG during lymphocytic choriomeningitis virus infection (Stolp et al., 2020) assuming similar motility characteristics of both cell types. T cells were observed to move in the SMG with an average speed of 6.8 μm/min, a top speed of 18 μm/min and were characterized with a low arrest coefficient (Stolp et al., 2020). Thus, these data indicate a fast motion with rare motility arrest and frequent direction changes.

Cell motility of CD4^+^ T cells was implemented in Morpheus using persistent motion, which is described by the persistence strength, sT, and the decay time 1T, defining the weight and the maintenance with which a cell pursues a specific target direction, respectively. The parameters were estimated using the FitMultiCell-pipeline (https://fitmulticell.gitlab.io) with simulations following 300 CD4^+^ T cells over a time period of 1h. Best estimates revealed a persistence strength of sT =180 and decay time of 1T =8.9 min assuming no deviation by chemotactic gradients (*CTX*=0). With these parameters, simulated T cell motility dynamics showed similar mean velocities and distribution of arrest coefficients, but constrained maximal velocities and higher straightness than observed in the experimental data (**Figure S7f**). These deviations are affected by the chosen temporal and spatial discretization of the simulation environment. Decreasing the time step size of the model would improve parameter fitting but substantially affects computational run time of the model. However, given the level of detail aimed for in our analyses, CD4^+^ T cell motility was sufficiently represented by the parameterization shown above.

#### CD4^+^ T cell stimulation, IFN*γ* dynamics and CXCL9/CXCL10 gradients

Within the SMG, CD4^+^ T cells can get stimulated to release IFN***γ,*** which inhibits viral replication within infected cells. In our simulations we distinguish between two scenarios of CD4^+^ T cells stimulation: (i) without and (ii) with the consideration of antigen presenting cells (APC). In a scenario without the consideration of APC, CD4^+^ T cells are stimulated by the direct contact with antigen, i.e., infected cells. The probabilities of endogenous and M25 CD4^+^ T cells to get stimulated, *p*_se_ = 1 − exp(-λ_se_) and *p*_sm_ = 1 − exp(-λ_sm_), respectively, were adjusted using previous observations from the literature(Verma et al., 2016). Assessing the percentage of IFN_γ_ secreting CD4^+^ T cells at 8 days post MCMV infection within various organs, Verma et al. (Verma et al., 2016) measured that 8 to 15% of CD4^+^ T cells, and 15-30% of epitope-specific cells were IFN_γ_^+^. To broadly reflect these dynamics, parameters λ_se_= 0.05 h^-1^ and λ_sm_ =0.15 h^-1^ were chosen. With this parameterization, we observed that 14.4 ± 5.17 % and 29.1 ± 12.1 % (mean ± SD of 10 replicates) of CD4^+^ and M25 CD4^+^ T cells, respectively, were IFN_γ_^+^ within our simulations (e.g. Figure 7 in the main manuscript), which is at the upper end of the observed intervals.

##### APC related activation

In the second scenario, we considered the additional effect of APC on CD4^+^ T cell motility dynamics and stimulation. APCs are suggested to have a macrophage-like phenotype that present MCMV-antigen after phagocytosis of apoptotic remains of infected cells (Walton et al., 2011). Therefore, in our model APCs were simulated as an additional layer with grid sites *APCij* (**Figure S7a**). The death of infected cells can lead to local antigen presentation, i.e., *APCij* = 1, at the grid site corresponding to the location of the infected cell. As infected cells tend to be close to each other and one APC might take up signals from different infected cells in the surrounding, only one out of three infected cells would on average result in antigen presentation. To represent CXCR3 guided motion, APCs are assumed to produce a local gradient of CXCL9/CXCL10 signaling that attracts CD4^+^ T cells. Different parameters were tested to define the chemotactic strength *CTX*_APC_ at which CD4^+^ T cells will move along this gradient. With values of *CTX*_APC_ = 10 and *CTX*_APC_ = 100 resulting in rare or recruitment of almost all CD4^+^ T cells to the specific APC, *CTX*_APC_ was set to 50. Upon contact with an APC, CD4^+^ T cells are stimulated with the same probabilities *p*_se_ and *p*_sm_ as before.

##### IFN_γ_ release

As CD4^+^ T cell motility is observed to be temporarily reduced after contact to an antigen (Honda et al., 2014), CD4^+^ T cells will stop their movement upon stimulation for on average of ∼42h, which corresponds to the lifetime of infected cells. The stopping time is therefore randomly determined by ρ(kI =0.25, 8I =170). During their immobility, cells will secrete IFN_γ_ for a confined time period, which was set to 2.5 h according to previous observations (Gao et al., 2014). Similar as for the viral concentration and infection spread, the effective range of IFN_γ_ is regulated by the diffusion coefficient for IFN_γ_ *D*_IFN_, the secretion rate *ρ*_IFN_, as well as the clearance rate, *c*_IFN_ (Eq. (7)). Assessment of these parameters is difficult and previous reported estimates, as e.g. for *ρ*_IFN_, vary substantially dependent on the method and system analyzed (Gao et al., 2014; Liao et al., 2014). We therefore used different parameter combinations of IFN_γ_ production, diffusion and clearance to test the effect of varying locally confined IFN_γ_ concentrations on the observed dynamics. Based on the values for production, degradation and diffusion, we can determine an effective IFN_γ_ perimeter, *r*_IFN_(*x*), given by the radius around the cell membrane up to which an IFN_γ_ concentration is achieved that would be at least as high as the EDx, thus leading to a x%- chance of inhibiting MCMV replication within infected cells within this area (see Eq. (5) and (6)). The perimeter is determined using the steady-state of Eq. (7). Defining an effective diffusion coefficient for IFN_γ_ of 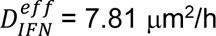, we adjusted the production and clearance rate to *ρ*_IFN_ = 25 ng cell^-1^h^-1^ and *c*_IFN_ = 0.15 h^-1^, respectively, to match our experimental observations **(Table A)**. With this parameterization, an individual CD4^+^ T cell would have an effective IFN_γ_ perimeter with a radius of *r*_IFN_(50) =10.77 μm, *r*_IFN_(25) = 18.97 μm, and *r*_IFN_(10) = 29.53 μm for a 50%, 25% and 10% chance, respectively, in inhibiting MCMV replication. This would correspond to an area of 364.42 μm^2^, 1130.97 μm^2^ or 2739.47 μm^2^ around the CD4^+^ T cell that would be protected by IFN_γ_ concentrations that are at least as high as the required ED50, ED25, or ED10, respectively. The impact of varying values of 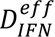, *ρ*_IFN_ and *c*_IFN_, and thus, varying effective radii for IFN_γ_ release on the progression of infection and protection was investigated.

#### Effect of IFN*γ* on viral replication

IFN_γ_ is assumed to suppress MCMV replication in the infected cells by late gene inhibition and was previously reported to be dose-dependent and reversible (Kropp et al., 2011; Lucin et al., 1994; Presti et al., 1998). In our model, the probability to inhibit viral replication in infected cells depends on the dose response relationship, *g(IFN)*, given in Eq. (6). The individual parameters, i.e., the maximal efficiency replication inhibition per time step, *y*_max_, the *ED*_SO_, and the Hill-coefficient *n* that characterize this dose-response relationship were determined using data from Lucin et al. (Lucin et al., 1994). In their experiments, Lucin et al. (Lucin et al., 1994) pretreated MEFs with different concentrations of IFN_γ_ for 48h, and determined the percentage of inhibition of viral load measured in pfu at 4 dpi. Even low concentrations of IFN_γ_ (<1 ng/ml) resulted in a reduction of replication of approximately 40%, and the effect quickly saturated with a maximum suppression of <60% at IFN_γ_ concentrations of >5 ng/ml. We fitted

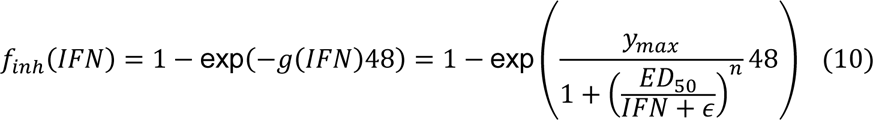

to these experimental data, with *f*_inh_(*IFN*) representing the percentage of inhibition dependent on the concentration of IFN_γ_, *IFN*, after 48h of treatment. Best fits and 95%-prediction intervals are shown in **Figure S7g** with best estimates given by *y*_max_= 1.76×10^-2^ h^-1^ (1.61×10^-2^, 2.0 ×10^-2^), *ED*_SO_= 0.41 ng/ml (0.27, 0.77), and *n* = 0.64 (0.51, 0.77) with numbers in brackets indicating 95%-confidence intervals of parameter estimates (see also **Table A**).

#### General system settings

The grid size, as well as simulated time step sizes had to be chosen in a trade-off between the required system size, level of detail and simulation duration. As mentioned above, our simulated grid size covered an area of 800×800 μm^2^ with 1 pixel (=lattice site) representing 2 μm in real space (**Table A**). This increased scaling allows the simulation of areas that are that large while still being able to determine spatiotemporal dynamics on a cellular level with a reasonable resolution. In addition, we used periodic boundary conditions and von-Neumann neighborhoods. Simulations were run based on a time step size of 1 hour. The duration of a MonteCarlo-Simulation time step (,i.e., MCS-duration), which determines the frequency of copy attempts of the CPM per simulation time step, was chosen to allow for realistic CD4^+^ T cell motility dynamics while retaining feasible simulations times. We used a MCS-duration of 0.0015, which resulted in > 600 MCS per time step (= 1 h). All simulations were run over 60 days with one run taking around 48 h on the high-performance computing cluster bwHPC MLS-WiSO. At the start of each simulation to initiate infection, two actively replicating MCMV infected cells were randomly placed within the simulated grid. In total, 7 different underlying duct structures were analyzed.

#### Parameter fitting and estimation procedures

Parameter fitting for ODE-models describing the dynamics of intracellular viral replication, CD4^+^ T cell influx, and IFN_γ_ efficacy were performed using Maximum-Likelihood approaches based on the R-packages FME (Soetaert. K.; Petzoldt, 2019), deSolve, as well as the *nls*-function from the base-package. Parameter identifiability and 95%-confidence intervals of estimates were determined by profile likelihood analysis (Raue et al., 2009) using the R-package ProfilIroning (https://github.com/GabelHub/ProfileIroning), or the function *nls.profile* from the base-package (IFN_γ_-dose response).

Parameter estimation for individual CD4^+^ T cell velocity within the CPM was performed using the FitMultiCell pipeline (https://fitmulticell.gitlab.io), which is based on combining Morpheus with the computational parallelization and high-performance approach pyABC to automatically adjust stochastic multi-scale models to experimental data. The pyABC workflow tests multiple parameter sets in parallel by subsequently minimizing distance measurements between experimental and simulated data. To this end, each simulation is evaluated using the R-Motilitylab package (http://www.motilitylab.net) to calculate the mean square displacement, the velocity, straightness and arrest coefficient of each cell. To account for differences in the measurement scales of these variables, the coefficient of variance is considered within the fitting procedure by

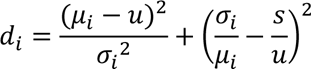

defining the calculated distance for the respective quantity *i*. Hereby, μ_i_ and σ_i_, are the mean and standard deviation of the experimental data, and *u* and *s* the corresponding values for the simulated data.

#### Data and Code availability

The individual codes and files describing the model structure to be run in Morpheus are made publicly available via GitHub (*Github-Link will be added here*).

---

**Table A:** General systems settings for the cellular Potts models and parameterization of the individual processes of cellular turnover, intracellular viral replication, viral clearance and diffusion, as well as CD4^+^ T cell dynamics and IFN_γ_-production as used within the different simulations. Numbers marked with (*), (#) or (†) indicate parameters that were estimated, adjusted or set, respectively, based on experimental observations and data from the literature. Sources are given within the text. (¶) indicates values that were determined based on data presented within this manuscript. Numbers in round-brackets indicate 95%-confidence intervals of estimates.

| Parameter | Description | Unit | Value |
| --- | --- | --- | --- |
| **General systems settings** |  |  |  |
| | Size of the grid | pixel<sup>2</sup> | 400 × 400 (1 px = 2 μm) |
|  | Time step duration | h | 1 |
|  | Duration of a MonteCarlo time step (MCS-duration) |  | 0.0015 = 5.4 s |
| | Epithelial cell area (uninfected) | μm<sup>2</sup> | 120.8 (¶) |
| | Epithelial cell area (infected) | μm<sup>2</sup> | 241.6 (¶) |
| | T cell area | μm<sup>2</sup> | 50 (¶) |
| **Epithelial cell dynamics** |  |  |  |
| $\lambda$ | max. proliferation rate of epithelial cells | h<sup>-1</sup> | 0.204 (#) |
| $N_{\max}$ | max. number of epithelial cells | cells | 2500 (¶) |
| $p_d$ | Probability for cell death of uninfected cells | | $\sim \Gamma(k_T = 40, \theta_T = 55)$ (†) |
| $p_\delta$ | Probability for cell death of infected cells | | $\sim \Gamma(k_I = 0.25, \theta_I = 170)$ (†) |
| **Intracellular viral replication** |  |  |  |
| $\alpha_R$ | viral net-replication rate | h<sup>-1</sup> | 0.41 (*)<br>(0.38, 0.49) |
| $R_{\max}$ | max. carrying capacity of intracellular virus per cell | gen. copy equiv. | 1060 (*)<br>(1020,1260) |
| $\rho_R$ | viral export rate | pfu/ gen. copy equiv. $h^{-1}$ | 0.03 (*)<br>(0.02,0.11) |
| $c_R$ | viral clearance rate (intracell.) | $h^{-1}$ | 0.45 (*)<br>(0.20,1.65) |
| **Extracellular viral concentration** |  |  |  |
| $c$ | viral clearance rate (extracell.) | $h^{-1}$ | 0.45 (*)<br>(0.20,1.65) |
| $D_V^{eff}$ | effective viral diffusion coefficient for the simulations | $\mu m^2 h^{-1}$ | 64 (#) |
| $\beta$ | Cell-free transmission rate | $pfu^{-1}h^{-1}$ | $1.5 \times 10^{-3}$ (#) |
| **CD4<sup>+</sup> T cell dynamics** |  |  |  |
| $\sigma_{in, end}$ | influx rate of endogen. CD4<sup>+</sup> T cells | $\times 10^{-2} h^{-1}$ | 0.17 (*,¶)<br>(0.07, 0.31) |
| $\sigma_{out, end}$ | loss rate of endogen. CD4<sup>+</sup> T cells | $\times 10^{-2} h^{-1}$ | 0.17 (*,¶)<br>(0.07, 0.31) |
| $\sigma_{in, M25}$ | influx rate of M25 CD4<sup>+</sup> T cells | $\times 10^{-2} h^{-1}$ | 0.9 (*,¶)<br>(0.40, 0.31) |
| $\sigma_{out, M25}$ | loss rate of M25 CD4<sup>+</sup> T cells | $\times 10^{-2} h^{-1}$ | 0.9 (*,¶)<br>(0.40, 1.40) |
| $E_{0, end}$ | Initial concentration of endogen. CD4<sup>+</sup> T cells | cells | 960 (*,¶)<br>(510, 2580) |
| $E_{0, M25}$ | Initial concentration of M25 CD4<sup>+</sup> T cells | cells | 20 (*,¶)<br>(10, 460) |
| $s_T$ | Persistence strength | | 180 (*) |
| $\tau_T$ | Decay time | min | 8.9 (*) |
| $CTX_{APC}$ | Chemotactic strength to follow CXCL9/CXCL10 gradients | | 50 (#) |
| $\lambda_{se}$ | stimulation rate of end. CD4<sup>+</sup> T cells | h<sup>-1</sup> | 0.05 (#) |
| $\lambda_{sm}$ | stimulation rate of M25 CD4<sup>+</sup> T cells | h<sup>-1</sup> | 0.15 (#) |
| **IFNγ release and efficacy** |  |  |  |
| $y_{max}$ | max. rate of replication inhibition by | $\times 10^{-2} \text{ h}^{-1}$ | 1.76 (*)<br>(1.61, 2.0) |
| $ED_{50}$ | effective dose | ng/ml | 0.41 (*)<br>(0.27, 0.77) |
| $n$ | Hill-coefficient for dose-response relationship | | 0.64 (*)<br>(0.51, 0.77) |
| $\rho_{IFN}$ | Production rate of IFNγ | ng h<sup>-1</sup> cell<sup>-1</sup> | 25 (#) |
| $c_{IFN}$ | Degradation rate of IFNγ | h<sup>-1</sup> | 0.15 (#) |
| $D_{IFN}^{eff}$ | effective diffusion coefficient of IFNγ for standard parameterization | $\mu\text{m}^2 \text{ h}^{-1}$ | 7.81 (#) |
| **CXCL9/CXCL10 dynamics** |  |  |  |
| $\rho_{CX9/10}$ | Release rate of CXCL9/10 | h<sup>-1</sup> | 0.05 (#) |
| $c_{CX9/10}$ | Degradation rate of CXCL9/10 | h<sup>-1</sup> | 0.04 (#) |
| $D_{CX9/10}$ | Diffusion coefficient of CXCL9/10 | $\mu\text{m}^2 \text{ h}^{-1}$ | 8.0 (#) |
| $CTX_{APC}$ | Chemotactic strength of CD4<sup>+</sup> T cells following CXCL9/10 gradient | | 50 (#) |

**Table B:** CD4^+^ T cell dynamics based as observed on the cross sections of SMG. The observed cell numbers for endogenous and M25 CD4^+^T cells at 8 dpi with the determined conversion factor for the simulated area of 800×800 μm^2^, as well as the transformed cell counts at 0, 8, 14 and 30 dpi are shown. The latter were used in the analysis for determining the net-influx and efflux rates based on Eq. (9) (see also **Figure S7e** and **Table A**). Observations were based on 4 cross sections with the mean ± SD shown.

| Transformation factor of cell counts based on cross section image analyses (n=4) at 8 dpi |  |  |
| --- | --- | --- |
|  | Endogen. CD4<sup>+</sup> T cells | M25 CD4<sup>+</sup> T cells |
| Mean cell count (1×1 cm<sup>2</sup>) | 509.88 | 10.54 |
| Mean # cells (800×800 μm<sup>2</sup>) | 326.32 | 6.74 |
| Mean # cells (SMG, extrapolation) | 76501.06 | 30731.16 |
| Transformation factor | 234.43 | 4557.70 |
| Transformed cell counts for 800×800 μm<sup>2</sup> sections |  |  |
| Time (dpi) | Endogen. CD4<sup>+</sup> T cells | M25 CD4<sup>+</sup> T cells |
| 0 | 63 (± 29.6) | 0 (± 0) |
| 8 | 223 (± 101.1) | 6.4 (± 2.9) |
| 14 | 386 (± 91.1) | 3.48 (± 2.2) |
| 30 | 357 (± 92.5) | 0.3 (± 0.1) |

