## Supplementary figures and images for "Locally confined IFN_γ_ production by CD4^+^ T cells provides niches for murine cytomegalovirus replication in the salivary gland"

### Graphical abstract

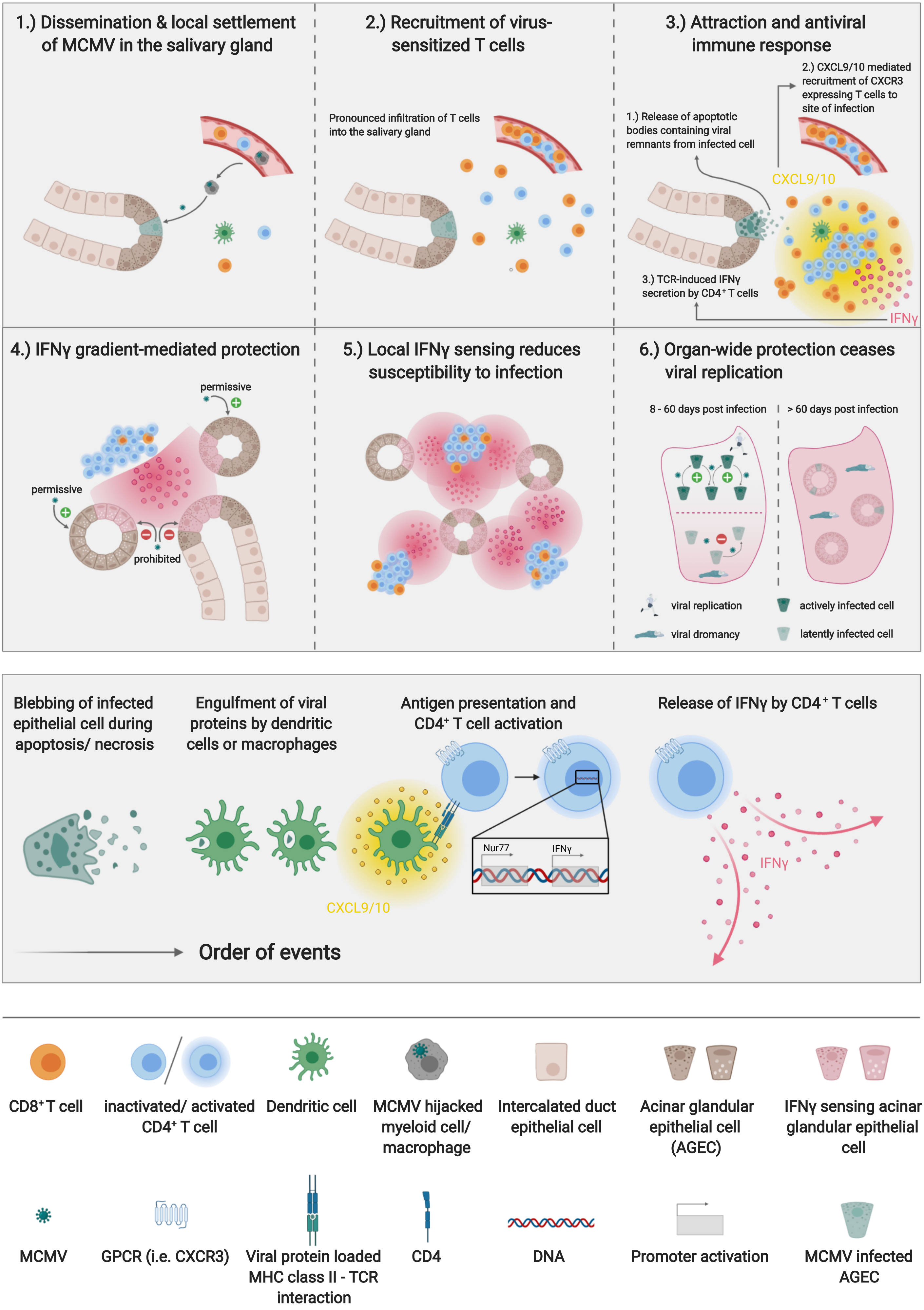
